# Developmental and adult striatal patterning of nociceptin ligand marks striosomal population with direct dopamine projections

**DOI:** 10.1101/2024.05.15.594426

**Authors:** Emily Hueske, Carrie Stine, Tomoko Yoshida, Jill R. Crittenden, Akshay Gupta, Joseph C. Johnson, Ananya S. Achanta, Johnny Loftus, Ara Mahar, Dan Hul, Jesus Azocar, Ryan J. Gray, Michael R. Bruchas, Ann M. Graybiel

**Affiliations:** McGovern Institute Brain Research and Department of Brain and Cognitive Sciences, Massachusetts Institute of Technology, Cambridge, MA 02139, USA; Center for the Neurobiology of Addiction, Pain and Emotion; Departments of Anesthesiology and Pharmacology, University of Washington, Seattle, WA 98105, USA; Molecular and Cellular Biology, University of Washington, Seattle, WA 98105, USA

**Keywords:** striatum, nociceptin, orphanin F/Q, striosome, dopamine, opioid, dendron bouquet

## Abstract

Circuit influences on the midbrain dopamine system are crucial to adaptive behavior and cognition. Recent developments in the study of neuropeptide systems have enabled high-resolution investigations of the intersection of neuromodulatory signals with basal ganglia circuitry, identifying the nociceptin/orphanin FQ (N/OFQ) endogenous opioid peptide system as a prospective regulator of striatal dopamine signaling. Using a prepronociceptin-Cre reporter mouse line, we characterized highly selective striosomal patterning of *Pnoc* mRNA expression in mouse dorsal striatum, reflecting early developmental expression of *Pnoc*. In the ventral striatum, Pnoc expression was was clustered across the nucleus accumbens core and medial shell, including in adult striatum. We found that Pnoc^tdTomato^ reporter cells largely comprise a population of dopamine receptor D1 (*Drd1*) expressing medium spiny projection neurons localized in dorsal striosomes, known to be unique among striatal projections neurons for their direct innervation of midbrain dopamine neurons. These findings provide new understanding of the intersection of the N/OFQ system among basal ganglia circuits with particular implications for developmental regulation or wiring of striatal-nigral circuits.

## 1 INTRODUCTION

The midbrain neurons of the substantia nigra influence motor and cognitive behaviors related to reinforcement learning, motivation, behavioral choice and movement initiation (da Silva et al. 2018, Hikosaka et al. 2014, Howe and Dombeck 2016, Schultz 2016). The circuits that innervate these dopamine neurons can bias, enhance or diminish these modulatory functions. A labyrinthine striosomal compartment of the striatum that winds through the surrounding matrix contains neurons unique among striatal projection neurons (SPNs) in their direct innervation of midbrain dopamine-containing neurons (Crittenden et al. 2016, Matsushima and Graybiel 2020, Watabe-Uchida et al. 2012). Many of their axons are known to innervate dendritic bundles of the dopamine neurons of ventral tier substantia nigra pars compacta (vSNc) and posterior cell cluster to form the striosome-dendron bouquets (Matsushima and Graybiel 2020). This striosome-to-dopamine dendron bouquet input can shut down activity of the dopamine neurons, with a following excitatory rebound (Evans et al. 2020). Here, we report that among these bouquet-innervating striosomal neurons there is a population selectively defined by the developmental expression of prepronociceptin (*Pnoc*), a gene encoding the opioid-like ligand nociceptin/orphanin FQ (N/OFQ) associated with pain, stress and reinforcement-related processing by way of its dampening effect on dopamine (Flau et al. 2002, Norton et al. 2002, Parker et al. 2019, Zheng et al. 2002).

Nociceptin/orphanin FQ (N/OFQ) is an opioid neuropeptide originally named for its induction of hyperalgesia (Meunier et al. 1995, Reinscheid et al. 1995). Its functions have since been shown to extend beyond pain modulation to influence motivation, stress, hormone regulation, learning and memory, mood regulation, feeding and substance use disorder (Jenck et al. 1997, Kash et al. 2015, Koob 2008, Mogil et al. 1996, Parker et al. 2019, D’Oliveira da Silva et al. 2023). It has also recently been considered as a treatment candidate for major depressive disorder (Der-Avakian et al. 2017, Gavioli and Calo’ 2013, Post et al. 2016, Ubaldi et al. 2021, Witkin et al. 2019). In the central nervous system (CNS), N/OFQ acts as a ligand for the N/OFQ opioid peptide (NOP) receptor, a G-protein coupled receptor that is widely distributed throughout the brain, including within the striatum (Mollereau and Mouledous 2000). In contrast to its receptor’s broad expression, N/OFQ ligand expression is more restricted and primarily localized in regions related to stress, feeding, and dopamine regulation (Neal et al. 1999, Norton et al. 2002, Ubaldi et al. 2021), including the central amygdala (CeA), hypothalamus and bed nucleus of stria terminalis (Hardaway et al. 2019, Hernandez et al. 2021, Kash et al. 2015, Smith et al. 2020), as well as in a population of dopamine-projecting neurons of the paranigral VTA neurons that locally modulate VTA dopamine neurons (Parker et al. 2019).

There is a strong precedent for the importance of opioid signaling in the striatum, with extensive literature pointing to numerous striatal functions of other endogenous opioid systems, most notably mu and kappa, in the coordination of key reward and feeding behaviors ((Bodnar et al. 1995, Ragnauth et al. 2000, Castro and Berridge 2014, Castro and Bruchas 2019, Castro et al. 2021). In addition, dysregulated striatal opioid signaling can contribute to a wide range of neurological disorders including mood disorders, substance use disorder, Huntington’s disease, Parkinson’s disease, and schizophrenia among others (Sgroi and Tonini 2018, Cui et al. 2014b, Banghart et al. 2015, Margolis et al. 2023, Morigaki et al. 2020, Kennedy et al. 2006, Canales and Graybiel 2000, Clark and Abi-Dargham 2019), in part owing to the distinct patterns of expression exhibited by different opioid receptors and peptides throughout striatum, including differential expression in striosome and matrix compartments (Crittenden and Graybiel 2011, Graybiel 1990, Brimblecombe and Cragg 2017, Tajima and Fukuda 2013). Although *Pnoc*-expressing neurons can be found throughout the striatum, their organization within striatal neuronal types and their anatomical intersection with dopamine circuitry across striatal subregions have remained largely unknown, especially in comparison to the other striatal opioid systems.

Despite this, several studies have implied functional significance for the N/OFQ system in both the dorsal striatum and nucleus accumbens. Studies of dorsal striatal dopamine release following activation or blockade of NOP receptors in SNc highlighted bidirectional modulation of dopamine (Marti et al. 2004, Murphy and Maidment 1999, Olianas et al. 2008) In the nucleus accumbens, N/OFQ infusion has been shown to reduce dopamine release (Koizumi et al. 2004, Murphy and Maidment 1999), and inhibit dopamine synthesis at DA nerve terminals in the NAc by reducing tyrosine hydroxylase phosphorylation (Olianas et al. 2008). With regard to behavioral function, N/OFQ infusion into the NAc core of rats reduces their motivation to pursue high-effort palatable food when a less palatable option is freely available (Wilson et al. 2024). Intracerebroventricular injection of N/OFQ also attenuates cocaine-induced increases to both locomotion and extracellular dopamine in the NAc (Lutfy et al. 2001). More broadly, N/OFQ has been implicated in the modulation of reward-related behaviors in studies showing that NOP receptor activation in the nucleus accumbens can reduce the rewarding effects of drugs of abuse such as cocaine (Bebawy et al. 2010, Caputi et al. 2014, Lutfy et al. 2002, Marquez et al. 2008, Murphy and Maidment 1999, Romualdi et al. 2007, Sakoori and Murphy 2004, Vazquez-DeRose et al. 2013). These findings suggest that N/OFQ signaling in the nucleus accumbens may be involved in approach-avoidance behavior and regulating the reinforcing properties of drugs.

While there is evidence of N/OFQ’s modulatory effects on striatal dopamine, little is known about the organization of striatal circuits subserving N/OFQ signaling and how they intersect with the midbrain dopamine system. This organization may be critical to N/OFQ’s influence on approach-avoidance and reward behaviors since several studies have demonstrated diversity in function of opioid peptide/receptor systems specific to their location within different striatal subregions (Resendez et al. 2013, Castro and Berridge 2014, Cui et al. 2014b, Al-Hasani et al. 2015, Tejeda et al. 2017, Castro et al. 2021). Notably, microstimulation of the other opioid receptors, mu, delta, and kappa, points to highly localized functional opioid ‘hotspots’ even within a single subregion of the nucleus accumbens (Castro and Berridge 2014).

In the current study, we investigated the spatial and molecular expression pattern of the N/OFQ ligand prepronociceptin (*Pnoc*) across the striatum. To visualize *Pnoc*+ neurons, we crossed mice with prepronociceptin-driven expression of Cre recombinase (*Pnoc-Cre*) with the Cre-dependent Ai14 (tdTomato) reporter line to induce a lasting tdTomato fluorescent label in Pnoc+ neurons (Pnoc^tdTomato^). We found that Pnoc^tdTomato^ neurons exhibit highly selective striosomal patterning in the dorsal striatum, and within the ventral striatum found distinctly concentrated Pnoc^tdTomato^ cell-clusters in the core and medial shell, but not in the lateral shell. In the dorsal striatum, Pnoc^tdTomato^ neurons predominantly colocalized with *Drd1*-expressing SPNs within striosomes, along with a smaller largely non-striosomal population of Pnoc^tdTomato^ interneurons co-expressing somatostatin (SST) and/or neuropeptide Y (NPY). In the ventral striatum, *Drd1* was similarly the the leading marker found co-expressed with Pnoc^tdTomato^ neurons, however a higher proportion of Pnoc^tdTomato^ neurons did not overlap with either *Drd1* or *Drd2* expression, particularly in the NAc shell. We also identified a population of Pnoc^tdTomato^ interneurons co-expressing NPY that is enriched in the nucleus accumbens relative to the dorsal striatum.

In contrast to the Pnoc^tdTomato^ patterning labeled by the reporter line, we found low detectable expression of *Pnoc* mRNA in the dorsal striatum of adult mice, in contrast to the ventral striatum where *Pnoc* mRNA expression persisted into adulthood. In addition, we observed robust striosomal patterned *Pnoc* mRNA expression in neonatal dorsal striatal tissue that is reflective of the later Pnoc^tdTomato^ patterning. Together, these findings are indicative of early developmental peptide expression that is likely eventually down-regulated. Finally, we found that fibers emanating from our identified striosomal Pnoc^tdTomato^ SPNs in the dorsal striatum forge a direct projection to dopamine-containing neurons of the substantia nigra pars compacta (SNc), intertwining their terminal arborizations with bundles of ventrally descending dendrites of nigral dopamine neurons in structures readily identifiable as striosome-dendron bouquets. Our findings presented here reveal the anatomical and molecular organization of *Pnoc*-expressing neurons across differ-Striatal nociceptin patterning ent striatal subregions and neuronal subpopulations, indicative of the structure that underlies N/OFQ’s functional interactions with striatal-nigral circuitry. They also highlight the dynamic expression throughout development of an opioid neuropeptide ligand known for its modulatory effects on behavior and dopamine function. These results suggest that *Pnoc* expression may be especially important in the early postnatal dorsal striosomal system, while its persistent expression in the ventral striatum points to the potential for this region’s influence on behaviors via N/OFQ signaling throughout development and into adulthood.

## 2 MATERIALS AND METHODS

### Mouse lines and husbandry

All animal procedures were approved by the Committee on Animal Care at the Massachusetts Institute of Technology, were performed in accordance with the U.S. National Research Council Guide for the Care and Use of Laboratory Animals Animals were maintained under conditions of constant temperature (25°C) and humidity (50%), and a 12:12 h light/dark cycle.The following mouse strains were used for behavioral, neural recording and histological experiments: *Pnoc-Cre* (Parker et al. 2019), Ai14 (Madisen et al. 2010) and LSL-Flpo (He et al. 2016). We crossed *Pnoc-Cre* mice on a C57BL/6J background (Jackson Laboratory) to the Cre-dependent Ai14 (tdTomato reporter line) or Cre-dependent LSL-Flpo line on FVB background (Taconic Biosciences) corrected for behavioral optimization. Consequently, F1 hybrid mice were produced to suit behavioral characterization (Menalled and Brunner 2014), avoiding the early hearing loss of the C57BL/6J mouse strain and poor vision of the FVB strain.

### Surgical preparations of stereotactic striatal injections

All surgeries were performed on adult mice aged 8-10 weeks. Animals were anesthetized with 5% isoflurane and maintained at 1.5-2% throughout the surgery. For intrastriatal delivery of viral vectors, mice were injected unilaterally with Flp-dependent membraneGFP and synaptophysin-mRuby constructs (AAV2/8 vector packaging hSyn-FDIO-mGFP-T2A-Synaptophysin-mRuby; Biohippo, Gaithersburg, MD) targeting the anterior dorsal striatum. All viruses were injected at a rate of 100 nL/min. Injection coordinates empirically optimized for F1 hybrid C57BL/6J x FVB mice were based on (Friedman et al. 2020) for ages <6 months: AP: +1.1 mm; ML: ± 1.7 mm; DV: 2.0 mm from brain surface.

### Chronic immobilization stress

Mice were placed in a Decapicone bag with an opening for the snout (model DC M200, Braintree Scientific) and partially immobilized for 1 hour per day for 14 consecutive days as previously described (Friedman et al. 2017). The bag was large enough not to impede breathing but small enough to limit mobility. The bags were taped across the back of the chest and taped shut around the tail so that the animals’ tails protruded. The mice were secured in an upright position to the bottom of an empty mouse cage. Mice were monitored every 5 min to ensure that breathing was not restricted and that they remained upright, unharmed, and fully contained.

### Chronic amphetamine

Drug injections were performed in a manner similar to previous reports (Crittenden et al. 2017). D-amphetamine hemisulfate salt (Sigma-Aldrich) was dissolved in 0.9% saline prior to use and injected i.p. at a dose of 7mg/kg. Mice were injected daily for 7 days at approximately 11am in their home cage, followed by 7 days without treatment and then a final ‘challenge’ day where a single dose of amphetamine (7 mg/kg) or saline was administered. The final challenge dose was followed by timed perfusion and brain tissue harvest 60 to 90 minutes after injection.

### Histological evaluation of brain tissue

Animals were transcardially perfused with a phosphate buffered saline solution (PBS) followed by 4% paraformaldehyde in PBS. Extracted brains were soaked in the paraformaldehyde solution for 24 hours and then maintained in 30% sucrose in PBS at 4°C for cryoprotection. After 24 hours, brains were frozen in dry ice and stored at −80°C. A sliding microtome was used to cut 30-μm thick sagittal or coronal sections from the brains of the adults, and 54μm-thick coronal sections from the brains of P1 to P20 mice. Sections were stored at 4°C in 0.02% sodium azide in 0.1 M PB ahead of preparation for immunofluorescence staining. For fluorescent in-situ hybridization (FISH), 10-μm thick coronal sections were cut from adult mouse brains and 18-μm thick coronal sections were cut from the P1 to P20 mouse brains. Sections were mounted on slide and stored at −80°C until use.

For immunofluorescence (IF) staining, sections were rinsed 3 times for 10 min in PBS with 0.2% Triton X-100 (Tx), then blocked with tyramide signal amplification (TSA) blocking solution (Akoya Bioscience FP1012) for 20 min at room temperature, and then incubated with primary antibodies (Table 1) diluted in TSA blocking solution for 48-72 hours at 4°C. Sections were again rinsed with PBS-Tx for 3 × 10 min, and were incubated with fluorescent conjugated secondary antibodies (Table 1) for 2 hours at room temperature. Sections were rinsed with PB for 10 minutes, then counterstained with DAPI solution diluted in PB for 2 minutes. After rinsing in PB for 3 times, sections were mounted with ProLong Gold and stored at 4°C until imaging. For FISH, RNAscope Multiplex Fluorescent Reagent Kit (Cat. 320850; Advanced Cell Diagnostics (ACD), Hayward, CA) and manufacturer’s protocol were used. Probes used in this study are shown in Table 2 with the combination of channel 1 to 3.

**TABLE 1.**
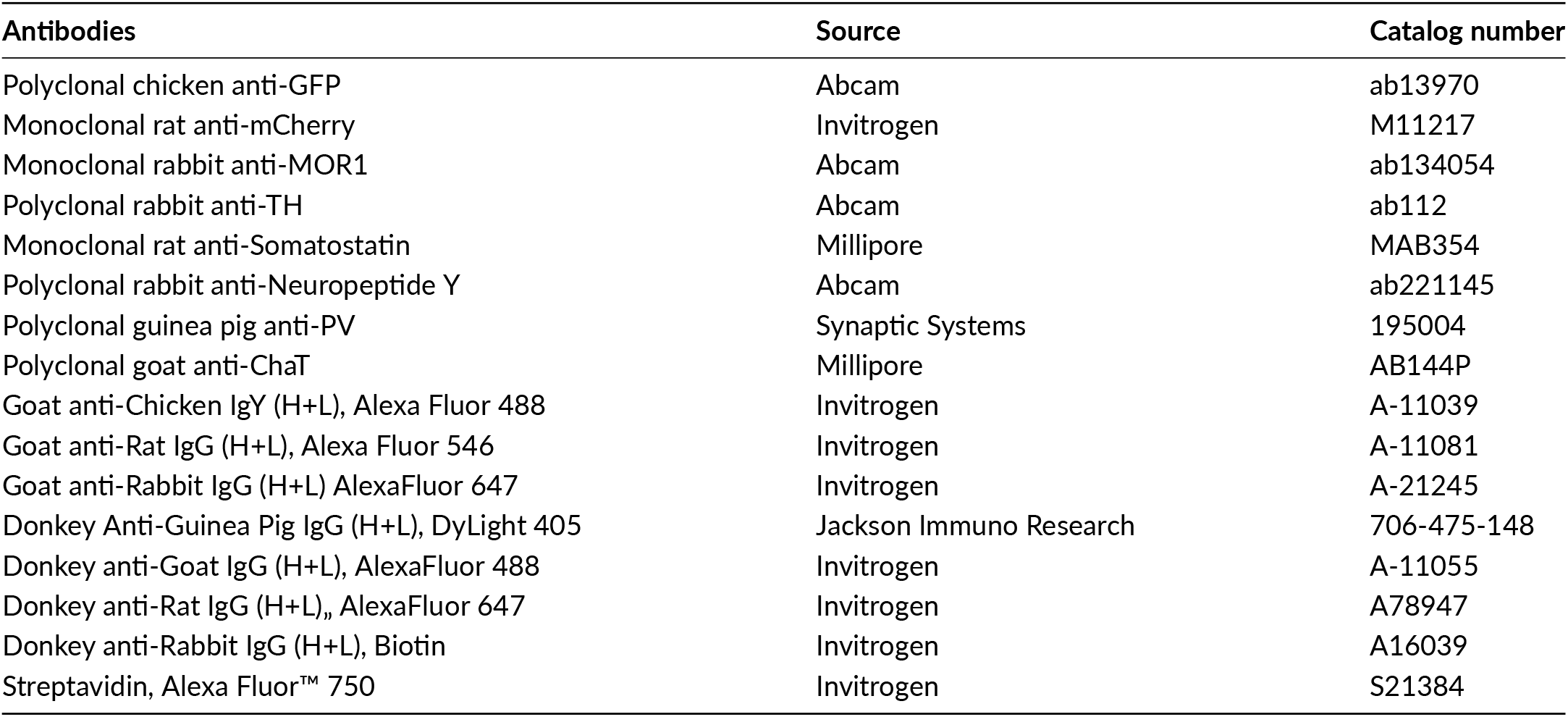
Antibodies used in this study.

| Antibodies | Source | Catalog number |
| --- | --- | --- |
| Polyclonal chicken anti-GFP | Abcam | ab13970 |
| Monoclonal rat anti-mCherry | Invitrogen | M11217 |
| Monoclonal rabbit anti-MOR1 | Abcam | ab134054 |
| Polyclonal rabbit anti-TH | Abcam | ab112 |
| Monoclonal rat anti-Somatostatin | Millipore | MAB354 |
| Polyclonal rabbit anti-Neuropeptide Y | Abcam | ab221145 |
| Polyclonal guinea pig anti-PV | Synaptic Systems | 195004 |
| Polyclonal goat anti-ChaT | Millipore | AB144P |
| Goat anti-Chicken IgY (H+L), Alexa Fluor 488 | Invitrogen | A-11039 |
| Goat anti-Rat IgG (H+L), Alexa Fluor 546 | Invitrogen | A-11081 |
| Goat anti-Rabbit IgG (H+L) AlexaFluor 647 | Invitrogen | A-21245 |
| Donkey Anti-Guinea Pig IgG (H+L), DyLight 405 | Jackson Immuno Research | 706-475-148 |
| Donkey anti-Goat IgG (H+L), AlexaFluor 488 | Invitrogen | A-11055 |
| Donkey anti-Rat IgG (H+L), AlexaFluor 647 | Invitrogen | A78947 |
| Donkey anti-Rabbit IgG (H+L), Biotin | Invitrogen | A16039 |
| Streptavidin, Alexa Fluor™ 750 | Invitrogen | S21384 |

**TABLE 2.** *In-situ* hybridyzation probes used in this study.

| Probe name | Source | Catalog number of channel 1 |
| --- | --- | --- |
| Mm-Drd1a | Advanced Cell Diagnostics | 406491 |
| Mm-Drd2 | Advanced Cell Diagnostics | 406501 |
| Mm-Pnoc | Advanced Cell Diagnostics | 437881 |
| tdTomato | Advanced Cell Diagnostics | 317041 |
| Mm-Oprl1 | Advanced Cell Diagnostics | 514301 |
| Mm-Oprm1 | Advanced Cell Diagnostics | 315841 |
| Mm-Lypd1 | Advanced Cell Diagnostics | 318361 |

For IF-FISH co-staining, RNase free blocking reagent and Codetection Target Retrieval Reagents from RNA-Protein Co-detection Ancillary Kit (Cat No. 323180, ACD) were used.

### Image analysis

Images were collected on a confocal TissueFAXs SL slide scanner (TissueGnostics) and were exported in tiff format. For the IF-FISH co-staining experiment, images were collected on the Epifluorescent TissueFAXs slide scanner (TissueGnostics) after IF staining (image set 1), then images were collected again following FISH (image set 2). These two image sets were exported in tiff format, then registered by the ImageJ plugin, StackRegJ (Stowers Institute for Medical Research). Registered images were then evaluated using ImageJ (FIJI), QuPath (Bankhead et al. 2017) or HALO image analysis software (IndicaLabs).

### Statistical analyses

Statistical analyses were performed using GraphPad Prism version 10.0.0 for Windows, GraphPad Software, Boston, Massachusetts USA, www.graphpad.com. All data are expressed as mean ± SEM.

## 3 RESULTS

### Pnoc^tdTomato^ cells in the striatum exhibit a patterned, clustered organization that corresponds with immuno-identified striosomes in the dorsal striatum

To make visible *Pnoc*-expressing neurons, we bred mice harboring *Pnoc-Cre* with a Cre-dependent “reporter line”, *loxP-Stop-loxP-tdTomato* (Ai14), such that offspring enduringly “report” red fluorescent protein, tdTomato, in cells that have shown expression of *Pnoc* (Pnoc^tdTomato^), whether current or previously. Examination of these Pnoc^tdTomato^ cells in the striatum demonstrated distinct spatial distributions across both dorsal and ventral regions (Figure 1). In the dorsal striatum, Pnoc^tdTomato^ cells were largely confined within striosomes, as identified by overlap with the known striosomal marker mu opioid receptor, MOR1 (Figure 1a-d). Discrete patterning was also evident in the ventral striatum, with Pnoc^tdTomato^ neurons mainly found predominantly in the dorsomedial part of the medial shell, within high-density clusters in the nucleus accumbens core and scattered sparsely and more homogeneously throughout the lateral shell (Figure 1a,e-g). Across the entire striatum, Pnoc^tdTomato^ neurons were distributed in similar proportions between the dorsal striatum, NAc core, and medial NAc shell, with a much smaller proportion found in the lateral NAc shell (Figure 1h). Pnoc^tdTomato^ neurons were consistently found in clusters that localized with striosomes along the dorsal striatum’s anterior to posterior axis, with only a scattered few found in the matrix (Figure 1i, Supplemental Figure 1). These findings demonstrate that *Pnoc* expressing neurons in the dorsal striatum are highly selective to the striosomal compartment and show clustered patterning in the ventral striatum.

**FIGURE 1.**
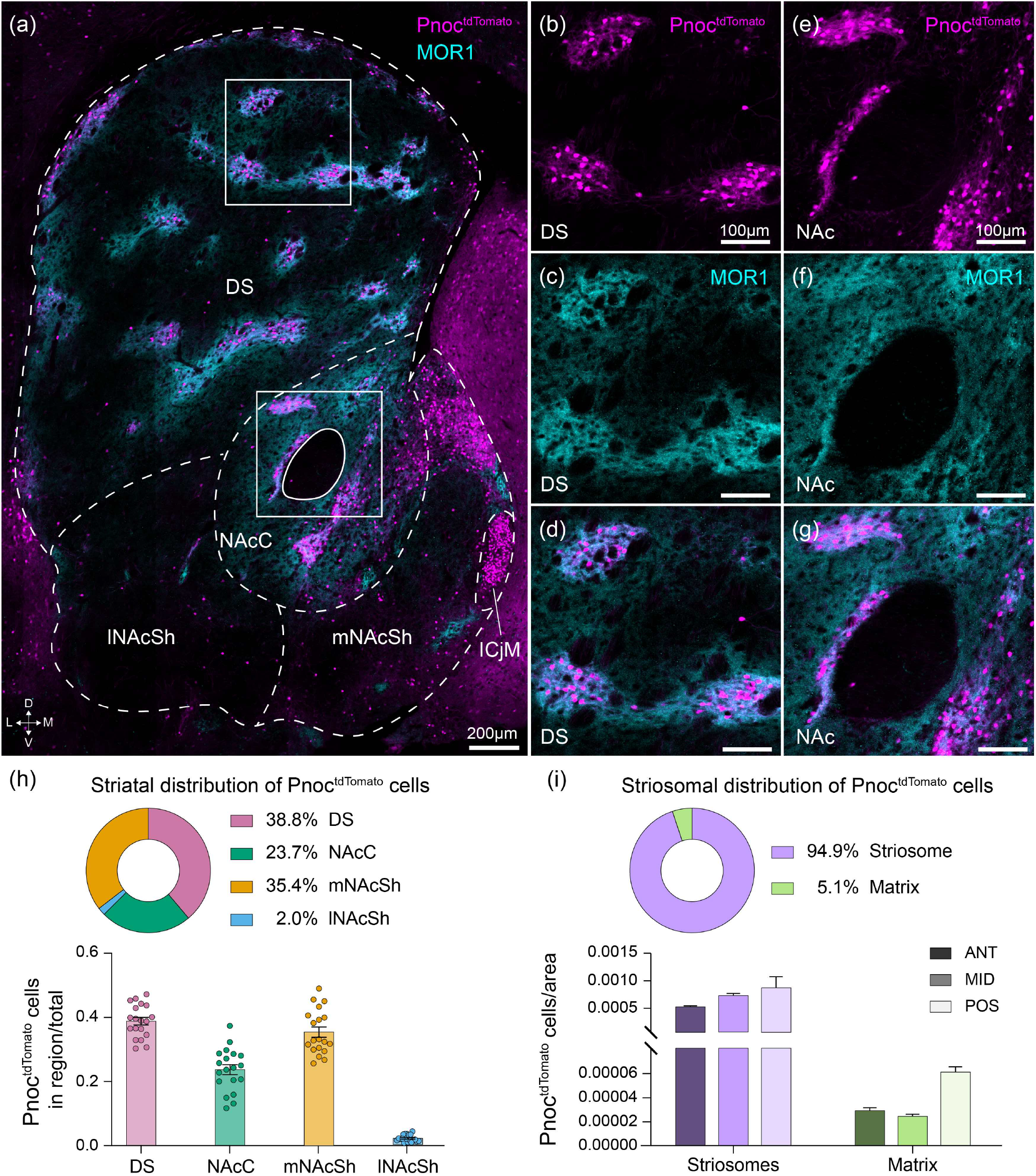
Pnoc^tdTomato^ reporter expression shows compartmental distribution across the striatum, and corresponds with striosomes in the dorsal striatum. (a) Composite image showing Pnoc^tdTomato^ (magenta) and striosomal marker MOR1 (cyan) in the striatum. (b-d) Higher magnification images of the top boxed region from panel (a) with Pnoc^tdTomato^ (b, magenta) and MOR1 (c, cyan) shown separately. Merged image (d) shows strong overlap between Pnoc^tdTomato^ neurons and Mor1 expression. (e-g) same as (b-d) but for lower boxed region in the ventral striatum from panel (a). Quantification of Pnoc^tdTomato^ spatial distribution across the dorsal and ventral (nucleus accumbens core, medial shell, and lateral shell) striatum. Quantification of Pnoc^tdTomato^ spatial localization in striosomes versus the matrix across the anterior to posterior axis in the dorsal striatum. DS = dorsal striatum; NAc = nucleus accumbens; NAcC = nucleus accumbens core; lNAcSh = lateral accumbens shell; mNAcSh = medial accumbens shell. ANT = anterior; MID = medial; POS = posterior.

### Striatal Pnoc^tdTomato^ neurons predominantly overlap with D1 SPNs, with greater heterogeneity among SPN subtypes in the ventral striatum

To further molecularly classify striatal *Pnoc*+ neurons, we evaluated Pnoc^tdTomato^ co-expression with markers for different MSN subtypes. We crossed our *Pnoc-Cre;LSL-tdTomato* reporter line with *Drd1-GFP* or *Drd2-GFP* reporter mice to simultaneously visualize Pnoc^tdTomato^ neurons labeled with either D1+ or D2+ neurons labeled with GFP (Figure 2). We found a significant colocalization of Pnoc^tdTomato^ with D1^GFP^-type SPNs in both dorsal (87.9 ± 0.7%) and ventral (64.9 ± 8.9%) striatum (Figure 2a-i), to the near exclusion of D2^GFP^ SPNs (Figure 2j-r). Due to the lack of full genetic penetrance characteristic of GFP reporter lines, we also quantified the SPN cell types by in situ hybridization (ISH) of *Drd1* and *Drd2*, which confirmed our initial findings (Figure 2s-u). The distribution of Pnoc^tdTomato^ neurons that co-localized with *Drd1, Drd2*, both, or neither, remained consistent across the dorsolateral, ventrolateral, and medial dorsal striatum (Supplemental Figure 2-1). By contrast, Pnoc^tdTomato^ SPN subpopulations in the ventral striatum appeared more heterogeneous, with just under half of Pnoc^tdTomato^ neurons in the lateral NAc shell (45.2 ± 10.6%) and almost a third in the medial shell (29.1 ± 9.1%) not co-localized with either *Drd1* or *Drd2* (Supplemental Figure 2-2). These results indicate that Pnoc^tdTomato^ labeling predominantly overlaps with *Drd1* expression across the dorsal and ventral striatum, suggesting that striatal *Pnoc* expression largely occurs within D1-type SPNs.

**FIGURE 2.**
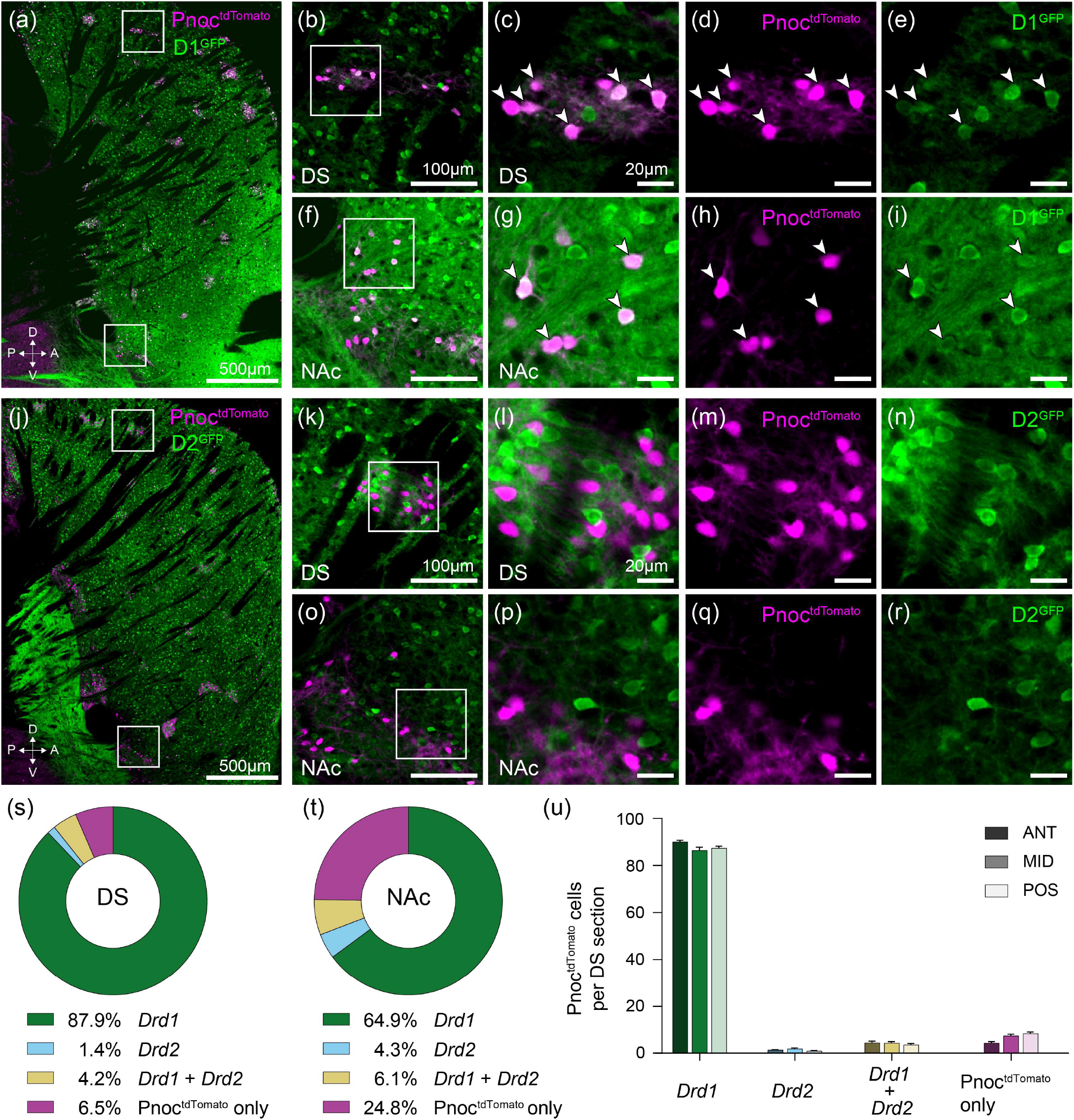
Pnoc^tdTomato^ expressing striatal neurons cluster with *Drd1* SPNs to near exclusion of *Drd2*. (a) Sagittal image of Pnoc^tdTomato^ cells (magenta) colocalizing with D1-GFP expressing SPNs (green) in dorsal (b-e) and ventral (f-i) striatum. (b) Higher magnification image of top boxed region from panel (a) in dorsal striatum. (c-e) Higher magnification images from inset in (b) of dorsal striatum. Arrowheads point to cells expressing markers for both Pnoc^tdTomato^ and D1-GFP. (f) Higher magnification image of bottom boxed region from panel (a) in ventral striatum (nucleus accumbens). (g-i) Higher magnification images from inset in (f) of ventral striatum. (j) Sagittal image of minimal overlap between Pnoc^tdTomato^ cells (magenta) and D2-GFP expression (green) in dorsal (k-n) and ventral (o-r) striatum (same as [b-e] and [f-i] but for D2-GFP expression). (s,t) Summary of IHC stained Pnoc^tdTomato^ reporter cells colocalizing with *in situ* hybridization-labeled *Drd1* and *Drd2* in SPNs in dorsal striatum (s) and nucleus accumbens (t). (u) Quantification of Pnoc^tdTomato^ cells on *Drd1* and *Drd2* labeled SPNs across the anterior to posterior axis in the dorsal striatum. DS = dorsal striatum; NAc = nucleus accumbens.

### A small, interneuronal population of Pnoc^tdTomato^ cells are marked by SST/NPY expression

We next asked whether any interneuronal populations of *Pnoc*+ cells in the striatum by staining for neuropeptide Y (NPY), somatostatin (SST), choline acetyltransferase (ChAT), and parvalbumin (PV) in combination with Pnoc^tdTomato^ labeling (Figure 3). The majority of striatal Pnoc^tdTomato^ neurons did not overlap with any interneuronal markers (dorsal striatum: 92.3 ± 0.5% Pnoc^tdTomato^ only; ventral striatum: 80.1 ± 4.0% Pnoc^tdTomato^ only), but a small dorsal striatal population of Pnoc^tdTomato^ neurons overlapped with NPY+/SST+ interneurons (5.2 ± 0.5%) often seen near the borders of striosomal Pnoc^tdTomato^ cell clusters, and a marginally larger subset in the ventral striatum overlapped with NPY+/SST-interneurons (17.7 ± 3.8%) (Figure 3a-e, k, l, Supplemental Figure 3). In both dorsal and ventral striatum, *Pnoc* reporter expression showed little to no overlap with PV+ or ChAT+ striatal interneurons (Figure 3f-j). These Pnoc^tdTomato^ interneuronal expression patterns did not detectably vary across the anterior to posterior striatal axis (Figure 3m). These multiple findings indicate that striatal *Pnoc*+ cells primarily overlap with D1-type SPNs and not striatal interneurons.

**FIGURE 3.**
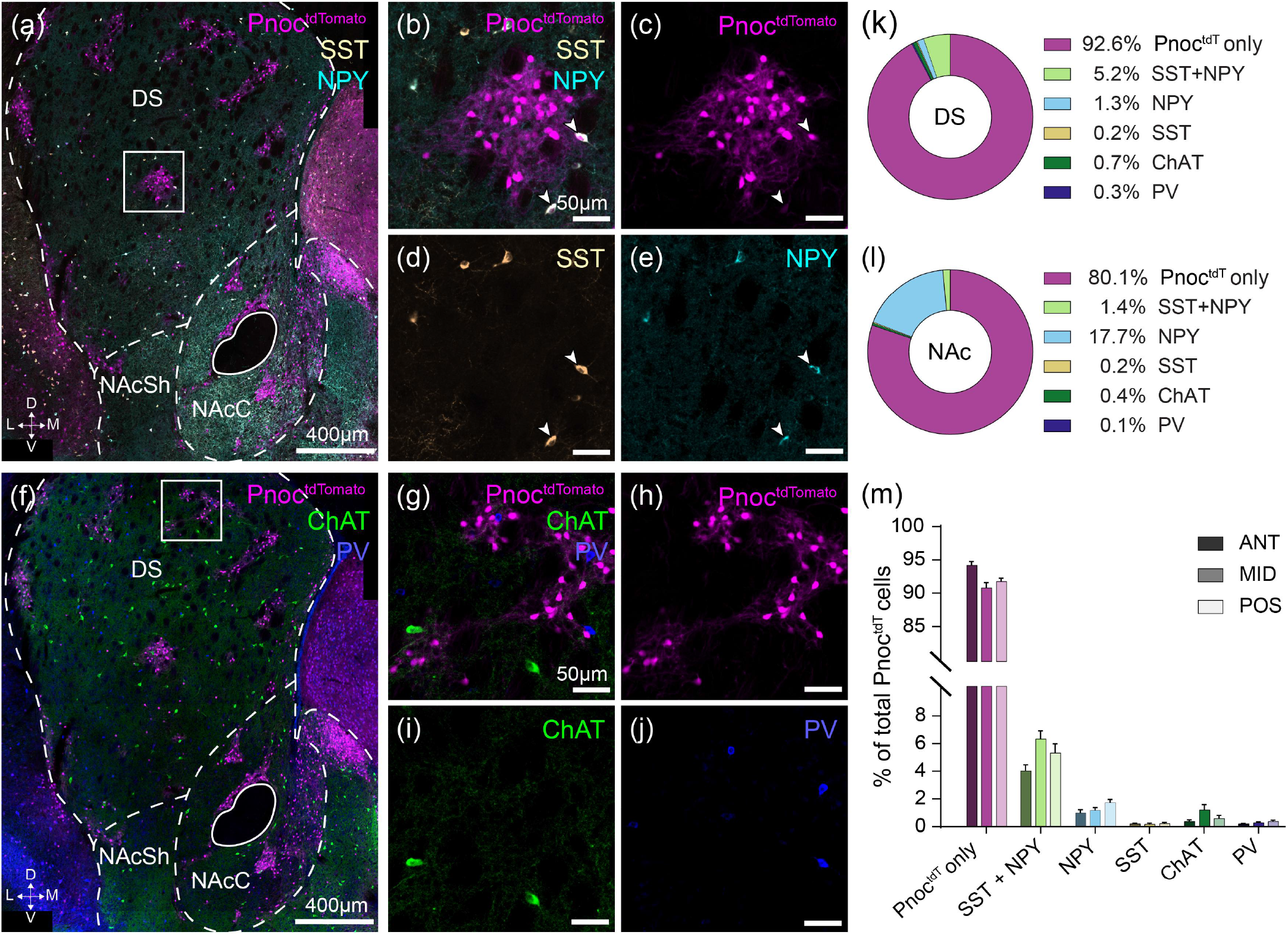
Pnoc^tdTomato^ reporter expression marks a population of NPY and SST co-expressing interneurons. (a) Coronal image of Pnoc^tdTomato^ cells (magenta), somatostatin (white, SST), and neuropeptide Y (cyan, NPY) expression in the striatum. Higher magnification images of inset from panel (a) for (b) merged and (c-e) individual channels showing Pnoc^tdTomato^ (c) colocalization with a subset of striatal interneurons co-expressing SST (d) and NPY (e). (f-j) Same organization as (a-e) for additional interneuron markers; Pnoc^tdTomato^ cells (magenta) show little colocalization with choline acetyltransferase (green, ChAT) or parvalbumin (blue, PV) expressing striatal interneurons. (k,l) Summary of quantified Pnoc^tdTomato^ cells colocalized with different interneuron markers in dorsal striatum (k) and nucleus accumbens (l). (m) Quantification of Pnoc^tdTomato^ cells colocalized with different interneuron markers across the anterior to posterior axis in the dorsal striatum. NAcC = nucleus accumbens core; lNAcSh = lateral accumbens shell; mNAcSh = medial accumbens shell.

### Expression of striatal *Pnoc* mRNA was low at adulthood and did not robustly increase following chronic exposure to either stress or to treatment with amphetamine

In contrast to the abundant *Pnoc*+ labeling found throughout the striatum using the tdTomato reporter line, we detected surprisingly little *Pnoc* mRNA expression in adult striatal tissue, particularly in the dorsal striatum (Figure 4; Supplemental Figure 4-1). Yet *Pnoc* mRNA expression was still frequently detectable in tdTomato labeled cells of the nucleus accumbens (Figure 4n-q, Supplemental Figure 4-1g-j). The lack of dorsostriatal Pnoc mRNA expression was especially striking since the *Pnoc* tdTomato reporter labeling in nearby septum and cortical regions did exhibit intense cellular *Pnoc* mRNA expression (Supplemental Figure 4-2k and 4-3k); the levels of *Pnoc* mRNA did not exceed negative control levels in dorsal striatal regions of reporter expression (Supplemental Figure 4-1c vs 4-1l).

**FIGURE 4.**
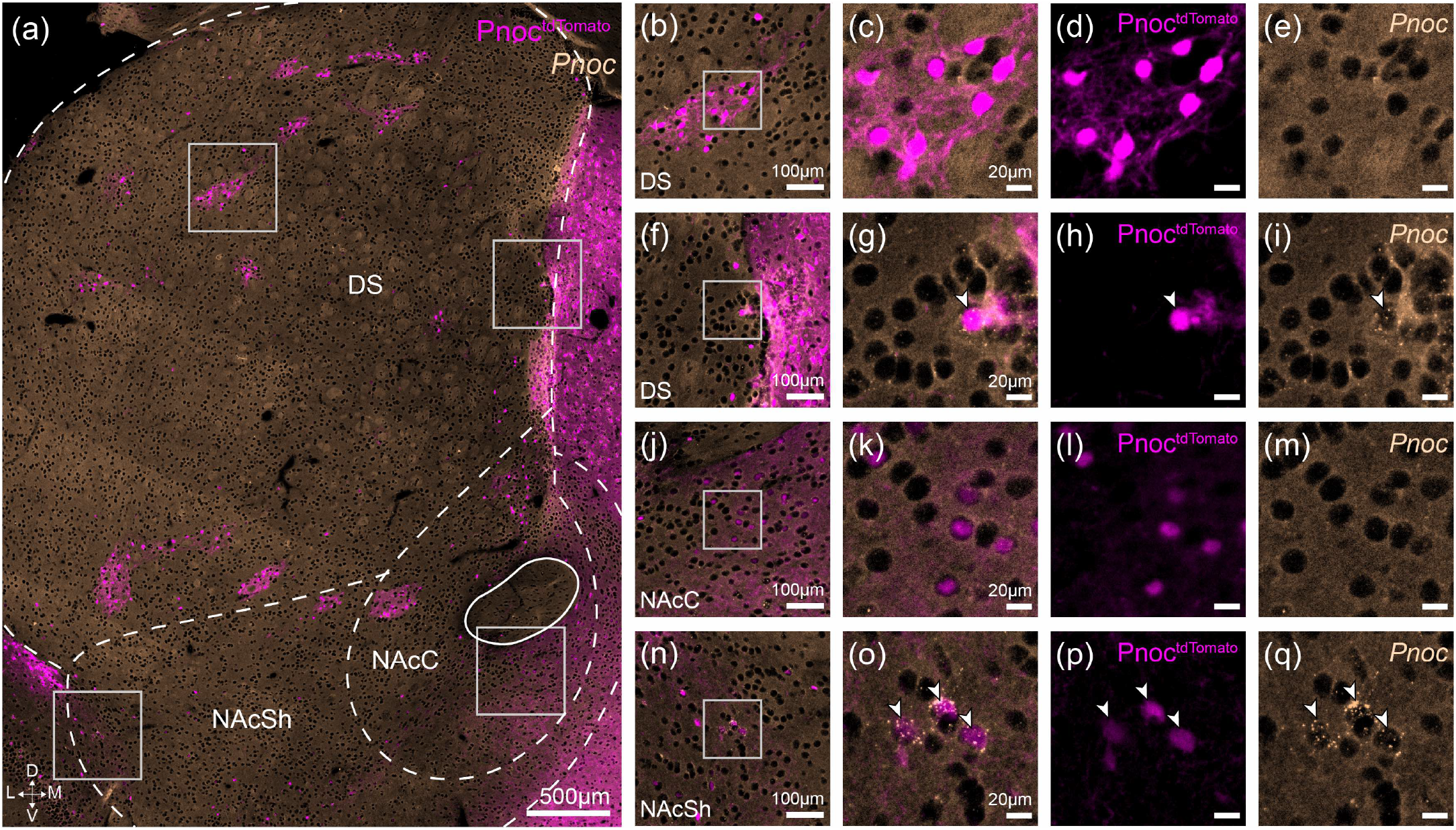
Adult expression of striatal *Pnoc* mRNA is sparse relative to reporter-labeled Pnoc^tdTomato^ cells. (a) Coronal image showing Pnoc^tdTomato^ cells (magenta) and *Pnoc* mRNA expression evaluated by ISH (yellow) in the adult striatum. Magnified insets from the dorsal striatum show Pnoc^tdTomato^ reporter-labeled cells that do not co-label (b-e) and do co-label (f-i) with adult *Pnoc* mRNA expression. Magnified insets from the nucleus accumbens show Pnoc^tdTomato^ cells that do not co-label (i-m) and do co-label (n-q) with adult *Pnoc* mRNA expression. DS = dorsal striatum; NAc = nucleus accumbens.

Given these puzzling findings, we asked whether striatal *Pnoc* expression could be induced in mature mice by activity-related or state-dependent changes. We first tested whether exposure to chronic stress would increase *Pnoc* expression in the dorsal striatum, given that midbrain N/OFQ receptor activation and some forms of chronic stress exposure have both been shown to disrupt striatal dopamine release (Imperato et al. 1992, Mangiavacchi et al. 2001). Daily chronic immobilization (1 hour/day for 14 days) produced no notable change in *Pnoc* mRNA expression as detected by ISH in the dorsal striatum (Supplemental Figure 4-2). We also tested whether exposure to a chronic dopamine agonist (amphetamine, 7 mg/kg for 7 days followed by a challenge dose on day 14 and tissue harvesting at 90 minutes post-dose) would induce robust adult *Pnoc* expression in the dorsal striatum, especially in striosomes, but again found very low *Pnoc* mRNA levels (Supplemental Figure 4-3). In summary, neither chronic stress nor chronic amphetamine exposure were viable avenues to induce strong striatal *Pnoc* mRNA expression in adult mice. Prompted by these observations, as well as by our observation of weak Cre-dependent recombination of reporter viruses injected into the mature striatum of *Pnoc*-Cre mice, we pursued alternative explanations behind the low *Pnoc* mRNA expression seen in adult tissue.

### *Pnoc* mRNA is developmentally expressed in striosomes

In *Pnoc-Cre;LSL-tdTomato* reporter animals, Pnoc^tdTomato^ labeling is triggered by the earliest sufficient expression of Cre-recombinase and will persist throughout the lifetime of the mouse. Given this, the discrepancy between the robust Pnoc^tdTomato^ reporter labeling and the dearth of adult *Pnoc* mRNA expression suggested that the *Pnoc* expression in the dorsal striatum could be developmentally regulated, with higher expression levels during embryonic and/or postnatal stages that are followed by a downregulation of expression as animals reach maturity. To investigate this possibility, we used ISH to detect *Pnoc* mRNA expression in the striatum of postnatal day 5 (P5) *Pnoc-Cre;LSL-tdTomato* reporter mice (Figure 5). In neonatal tissue, *Pnoc* mRNA expression did recapitulate the reporter-labeled pattern of Pnoc^tdTomato^ cells (Figure 5b-g). The neonatal *Pnoc* expression was largely in striosomes, consistent with the Pnoc^tdTomato^ expression pattern observed in adult mice (Supplemental Figure 5). These findings suggest that striosomal *Pnoc* expression is likely to be elevated during earlier developmental stages but is subsequently downregulated.

**FIGURE 5.**
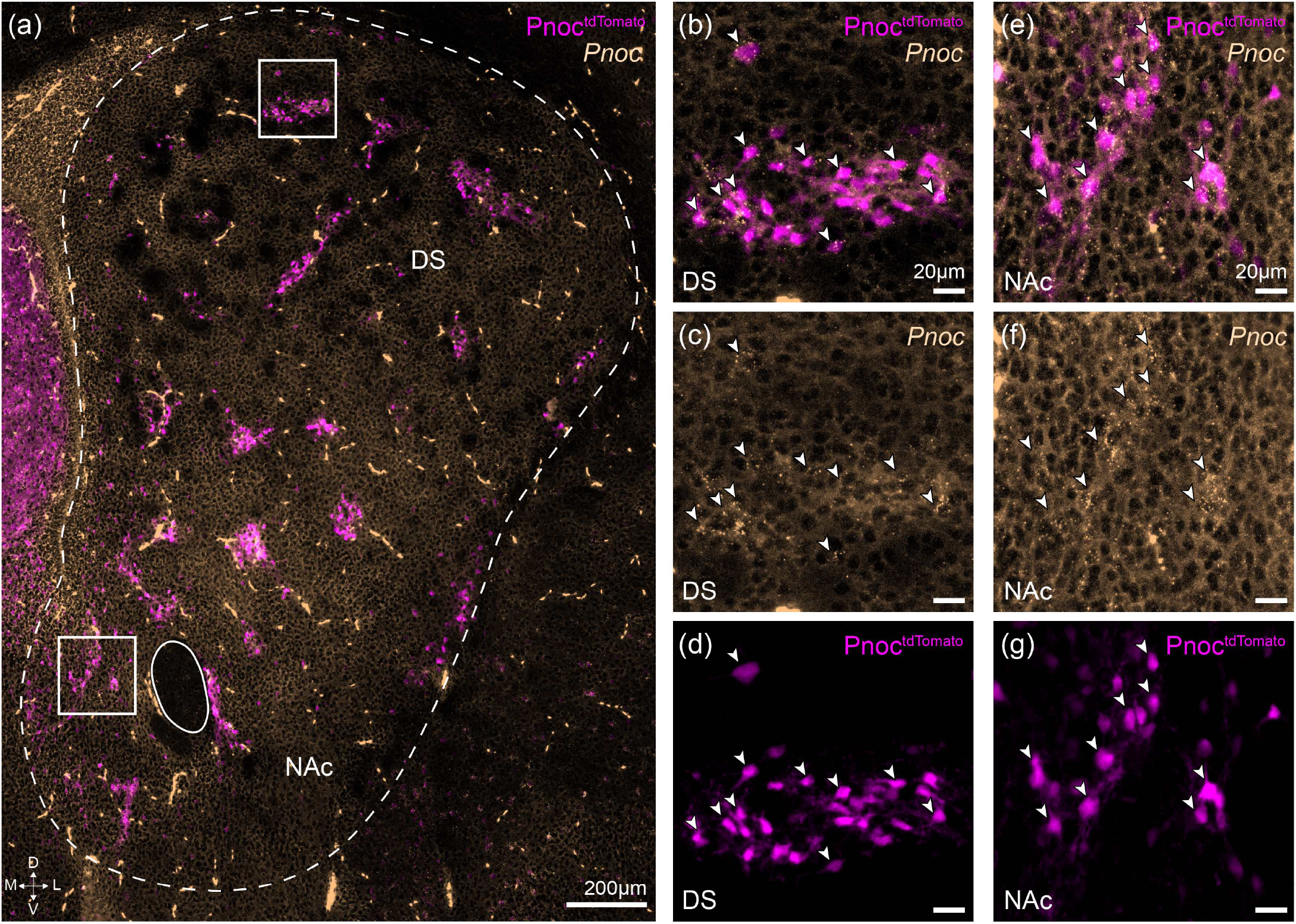
Neonatal expression of striatal *Pnoc* mRNA recapitulates reporter-labeled patterning in Pnoc^tdTomato^ cells. (a) Coronal image showing Pnoc^tdTomato^ reporter cells (magenta) and neonatal *Pnoc* mRNA expression evaluated by ISH (*Pnoc*, yellow) in the striatum of postnatal day 5 brain sections. Magnified insets from the dorsal striatum (b-d) and nucleus accumbens (e-g) show Pnoc^tdTomato^ reporter-labeled cells co-label with neonatal *Pnoc* mRNA expression. DS = dorsal striatum; NAc = nucleus accumbens.

### Pnoc^tdTomato^ neurons in striosomes directly project to dopamine neurons

Striosomal *Drd1* neurons are unique among striatal SPNs, or nearly so, in their direct innervation of midbrain dopamine neurons, in particular the ventral tier dopamine neurons of the SNc. The strong expression pattern in striosomal *Drd1* SPNs suggested that dorsal striatal Pnoc^tdTomato^ cells might correspond to this population. Examining midbrain sections of the Pnoc^tdTomato^ reporter mice, we observed specific innervation of the nigral striosome-dendron bouquets by the labeled axons of Pnoc^tdTomato^ cells both in the adult (Figure 6a-d) and in early P5 mouse tissue (Figure 6e-h). We then established that axons projecting specifically from striosomes account for these bundled fibers entwining themselves with dopamine-containing dendrons, rather than from another source, by injecting a Flp-dependent membraneGFP-expressing AAV vector into dorsal striatum of *Pnoc-Cre;loxP-Stop-loxP-FlpO* (Pnoc^FlpO^) mice (Figure 6i-n). Again the striosome-dendron bouquets were innervated by the population of striosomal neurons defined by early Pnoc expression. These findings demonstrate that *Pnoc*+ striatal neurons comprise a population of dorsal striatal dSPNs highly selective to the striosomal compartment, and that these *Pnoc*+ neurons forge, early in development, the specialized striatonigral circuit targeting striosome-dendron bouquets. Our results open the door for further investigation of the functional significance of this population and its ventral striatal counterpart, also showing clustered patterning of *Pnoc*+ but with persistent expression into adulthood.

**FIGURE 6.**
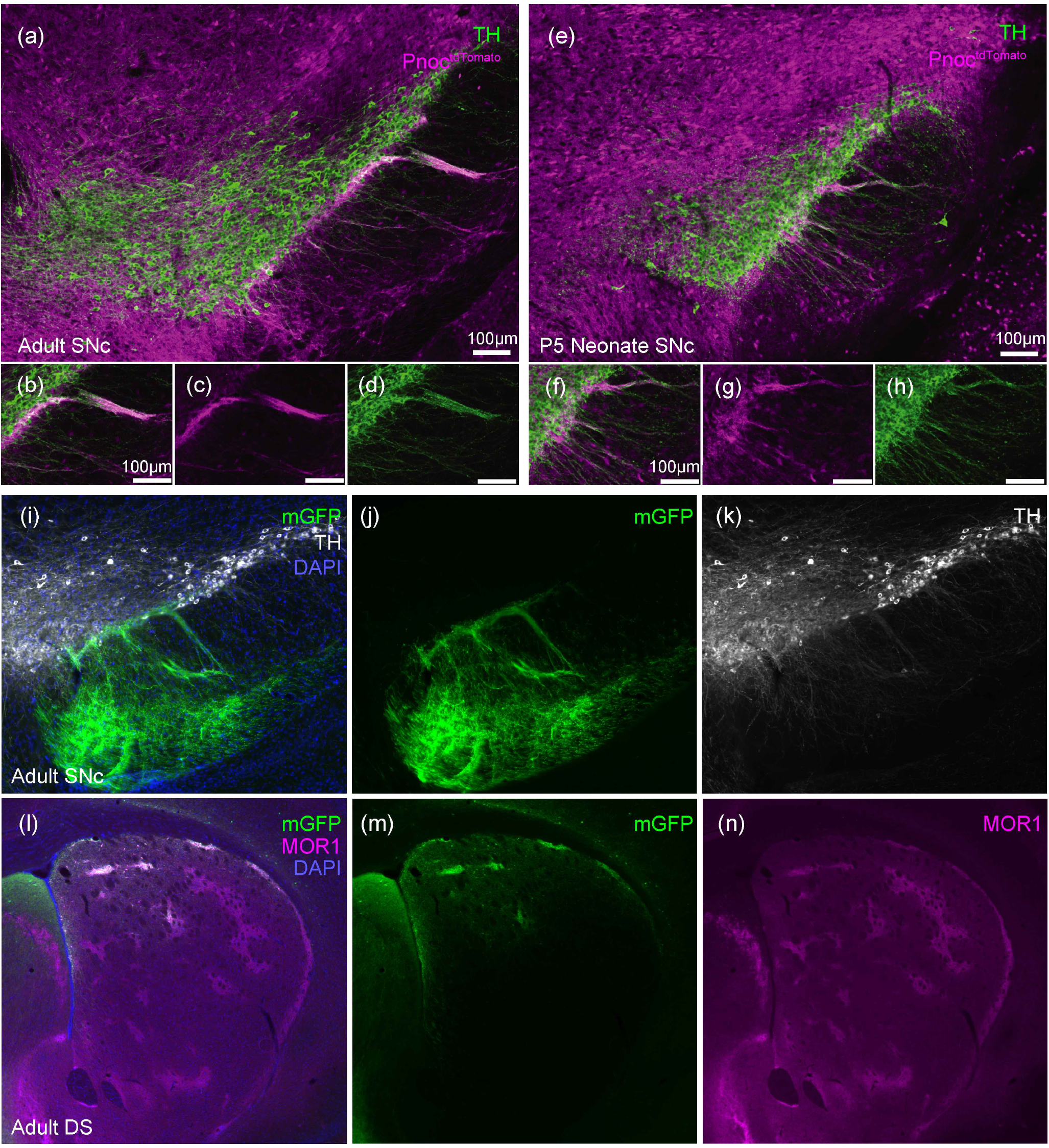
Striosomal fibers from Pnoc^tdTomato^ targeted neurons project to the striosome-dendron bouquet in SNc in both adult and neonate. In Pnoc^tdTomato^ reporter mice, tdTomato-labeled fibers (magenta) project to TH-labeled dopamine neurons (green) in adult (a-d) and neonate (e-h). Viral labeling of striosomal cells in P*noc-Cre;loxP-Stop-loxP-FlpO* mice with a Flp-dependent AAV expressing membrane-GFP (mGFP) exhibit projections to dopamine dendrons in SNc (i-k). mGFP labeled projections to dopamine neurons originate in virally-injected striosomes (l-n) Scale bars 100um.

## 4 DISCUSSION

The N/OFQ system has a demonstrated ability to disrupt the landscape of dopamine signaling with functional consequences relevant to stress and mood regulation, substance use disorder, and Parkinson’s disease among others. The striatal circuits and cell types that N/OFQ identifies so as to drive these effects have remained poorly defined. Here we lay the groundwork for investigating N/OFQ-mediated striatal contributions to these dopamine-related functions. In particular, we have demonstrated that striatal Pnoc^tdTomato^ cells reside with high selectivity in striosomes in the dorsal striatum and that they co-express *Drd1* but not *Drd2*, thereby comprising part of the unique striosomal direct pathway neurons that make direct projections with dopamine-containing neurons via striosome-dendron bouquet structures in the ventral tier SNc (Crittenden et al. 2016, Evans et al. 2020, Matsushima and Graybiel 2020, Watabe-Uchida et al. 2012).

For the ventral striatum, we highlight a newly identified population of molecularly heterogeneous *Pnoc* neurons that have distinct distributions across the nucleus accumbens shell and core and adjoining ventral territories. We also demonstrate that *Pnoc* expression in the ventral striatum is present from birth through adulthood. The distribution of Pnoc^tdTomato^ neurons in ventral striatum is more heterogeneous than dorsal Pnoc^tdTomato^ neurons both in cell identity and spatial distribution, but clear clustering is apparent in the accumbens core and medial shell, and across all subregions *Drd1* co-expression predominates. To determine whether these ventral striatal direct pathway neurons share a circuit motif of dopamine projection with their dorsal striosomal neighbors will require careful mapping of individual ventral striatal populations. Given the known importance of the ventral striatum for many of the functions for which N/OFQ has been implicated and in which dopamine is involved, we suggest that the afferent and efferent projections of these distinct ventral populations are an important next step alongside the targeting, including in the adult brain, of the NOP receptor ligand, N/OFQ, in pursuit of evaluating ventral striatal N/OFQ function.

We found Pnoc^tdTomato^ to be a superlative marker of striosomes across development and maturity, but that this patterning reflects developmentally transient expression of the Pnoc gene, which is downregulated by adulthood (Figures 4-5). Such transient developmental expression is reminiscent of a number of striatal genes that exhibit striosomal patterning early in development including FoxP2 (Enard 2011, Fong et al. 2018), Asc1 and Dlx1 (Kelly et al. 2018). Striosomal neurons are born during embryonic days 10.5-13.5 and lead the migration out into the striatal primordium ahead of the migration of neurons destined for the matrix compartment (Graybiel and Hickey 1982, Kelly et al. 2018, Lança et al. 1986, Matsushima and Graybiel 2020). In the course of development, striosomes and matrix exhibit different patterns of neurochemical and mRNA expression (Eblen and Graybiel 1995, Gokce et al. 2016, Graybiel and Hickey 1982, Kelly et al. 2018, Lança et al. 1986, Märtin et al. 2019, Matsushima and Graybiel 2020, Saunders et al. 2018) and develop distinct efferent and afferent patterns of connectivity. Striosomal neurons are more connected with limbic regions than are neurons of the matrix; they receive enriched inputs from limbic regions of cortex, bed nucleus of the stria terminalis, amygdala as well as providing the main source of striatal projections to midbrain dopamine neurons (Crittenden et al. 2016, Crittenden and Graybiel 2011, Graybiel and Matsushima 2023, Lee et al. 2024, Ragsdale and Graybiel 1988, Smith et al. 2016, Watabe-Uchida et al. 2012).

Disruption of striatonigral N/OFQ signaling during development could have profound long-term effects on dopamine-dependent behaviors, affecting not only movement-related activity but also aspects of mood, motivation, and possibly habit formation, including habitual aspects in the development of substance use disorder. N/OFQ shares amino acid sequence similarities with other opioid peptides, particularly dynorphin, which has high expression in parts of the striosomal system and, like other opioid mRNAs, for which expression is stimulated by cAMP or perturbations that raise cAMP levels. Studies following the discovery of N/OFQ showed that cAMP induces robust increases in expression of *Pnoc* mRNA, and that high levels of *Pnoc* transcript in undifferentiated neural cells is accompanied by neurite outgrowth including by transfection independent of cAMP (Saito et al. 1995 1996 1997, Zaveri et al. 2006). N/OFQ in turn activates the master transcriptional regulator, NFκB, which is important in gene regulation involved in synaptic plasticity, memory formation, cognition, development, and neuronal survival (Donica et al. 2011, Ahn et al. 2008) In addition, N/OFQ modulates immune factors such as TNFα and IL-6 (Fu et al. 2007) crucial to the formation and maintenance of neural circuit development and synaptic plasticity (Boulanger 2009). Our finding of robust *Pnoc* expression selectively in striosomal direct pathway neurons slated for dopaminergic interconnection opens the door for evaluation of how these molecular genetic and immune interactions unfold in a circuit of wide functional and pathological consequence. Still to be determined is the impact on the striatonigral circuitry of developmental perturbations that recruit these modulators of *Pnoc* expression.

The potential for development of circuit vulnerabilities and dysfunction through mis-wiring of striosomal neurons with the dopamine system is evident from the nature of those neurons with which striosomes form interconnections. Here, we show that the striosomal neurons that developmentally expressed *Pnoc* directly innervate vSNc dopamine neurons (Figure 6). The dopamine neurons of the vSNc, the recipient neurons of the striosomal-dendron bouquet axons, carry particular susceptibility to degeneration in Parkinson’s disease (Damier et al. 1999, Giguère et al. 2018, Kordower et al. 2013, Mendez et al. 2005, Pereira Luppi et al. 2021, Yamada et al. 1990). More widely, the cortico-striosome-dopamine circuit is implicated in mood regulation and dysregulation as striosomes are differentially implicated from surrounding matrix in reinforcement-related updating paradigms (Bloem et al. 2017 2022, Yoshizawa et al. 2018), and, in particular, in influencing cost-benefit conflict decision-making in both rodents (Friedman et al. 2015 2017 2020) as well as non-human primates (Amemori et al. 2018, Amemori and Graybiel 2012, Amemori et al. 2020 2021), findings that have been extended to human patient populations (Ironside et al. 2020, Pedersen et al. 2021, Rolle et al. 2022). These studies suggest circuits to study for a mechanistic understanding of how the cortico-striato-nigral circuit, and the nigro-striato-nigral loop, operate in dopamine regulation in the context of such disorders. As a result of its modulation of dopamine, the N/OFQ system has attracted interest as a therapeutic target for modulating dopamine in both Parkinson’s disease, substance use disorder, and mood regulation (Gavioli and Calo’ 2013, Lambert 2008, Post et al. 2016, Witkin et al. 2019, Zaveri 2011), including in alleviation of depressive symptoms resulting from developmental inflammatory insults arising from maternal immune activation (Medeiros et al. 2015). Our findings point to the value of further work on this developmental impact.

## Abbreviations

Pnoc: prepronociceptin
N/OFQ: nociceptin/orphanin FQ
NAc: nucleus accumbens
DS: dorsal striatum.

## AUTHOR CONTRIBUTIONS

EAH, CS, TY, JC, AMG, MRB conceived and designed experiments; TY, JL, AM collected the data; EAH, CS, TY, AG, JCJ, ASA, JL, RJG performed cell counting and image analysis; EAH, CS, TY, AG performed the statistical analyses; EAH, DH, JA performed surgical procedures and mouse husbandry; CS, EAH, AG, RJG prepared figures; EAH, CS, AMG, MRB wrote the paper.

## ACKNOWLEDGMENTS

We thank Sophia Robertson and Smitha Bhagavatula for assistance with histological cell counting. We thank Diego Pizzagalli for comments on the manuscript. This research was supported by NIH/NIMH P50 MH119467 (Project 3 to AMG; Project 4 to MRB), NIH/NIMH R01 MH060379 (AMG), NIH/NIMH F31 F31DA059438-01A1 (CAS), Saks-Kavanaugh Foundation (AMG), Hock E. Tan and K. Lisa Yang Center for Autism Research (AMG), Mallinckrodt Foundation (MRB), Simons Foundation grant to the Simons Center for the Social Brain at MIT (AMG).

## DATA AVAILABILITY

The data supporting this study’s findings are available upon request from the authors.

## SUPPORTING INFORMATION

Additional supporting information may be found in the online version of the article.

## SUPPLEMENTAL FIGURES

**SUPPLEMENTAL FIGURE 1.**
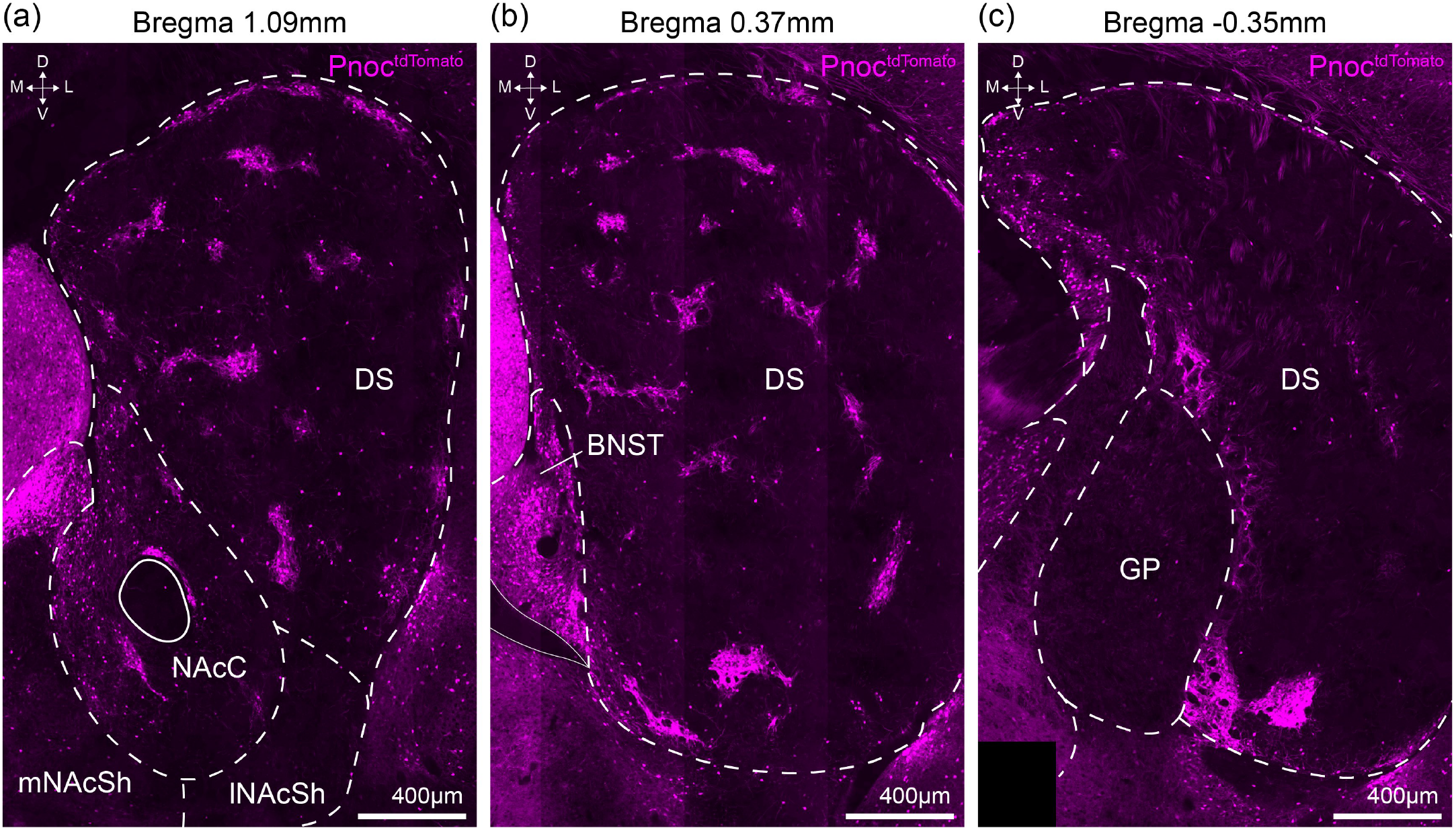
Pnoc^tdTomato^ striatal expression across the anterior to posterior axis. (a-c) Striatal Pnoc^tdTomato^ reporter expression (magenta) across the anterior (a, bregma = 1.09mm) medial (b, bregma = 0.37mm), and posterior (c, bregma = −0.35mm) striatum. DS = dorsal striatum, NAcC = nucleus accumbens core; mNAcSh = medial accumbens shell; lNAcSh = lateral accumbens shell; BNST = bed nucleus of the stria terminalis; GP = globus pallidus.

**SUPPLEMENTAL FIGURE 2. - 1.**
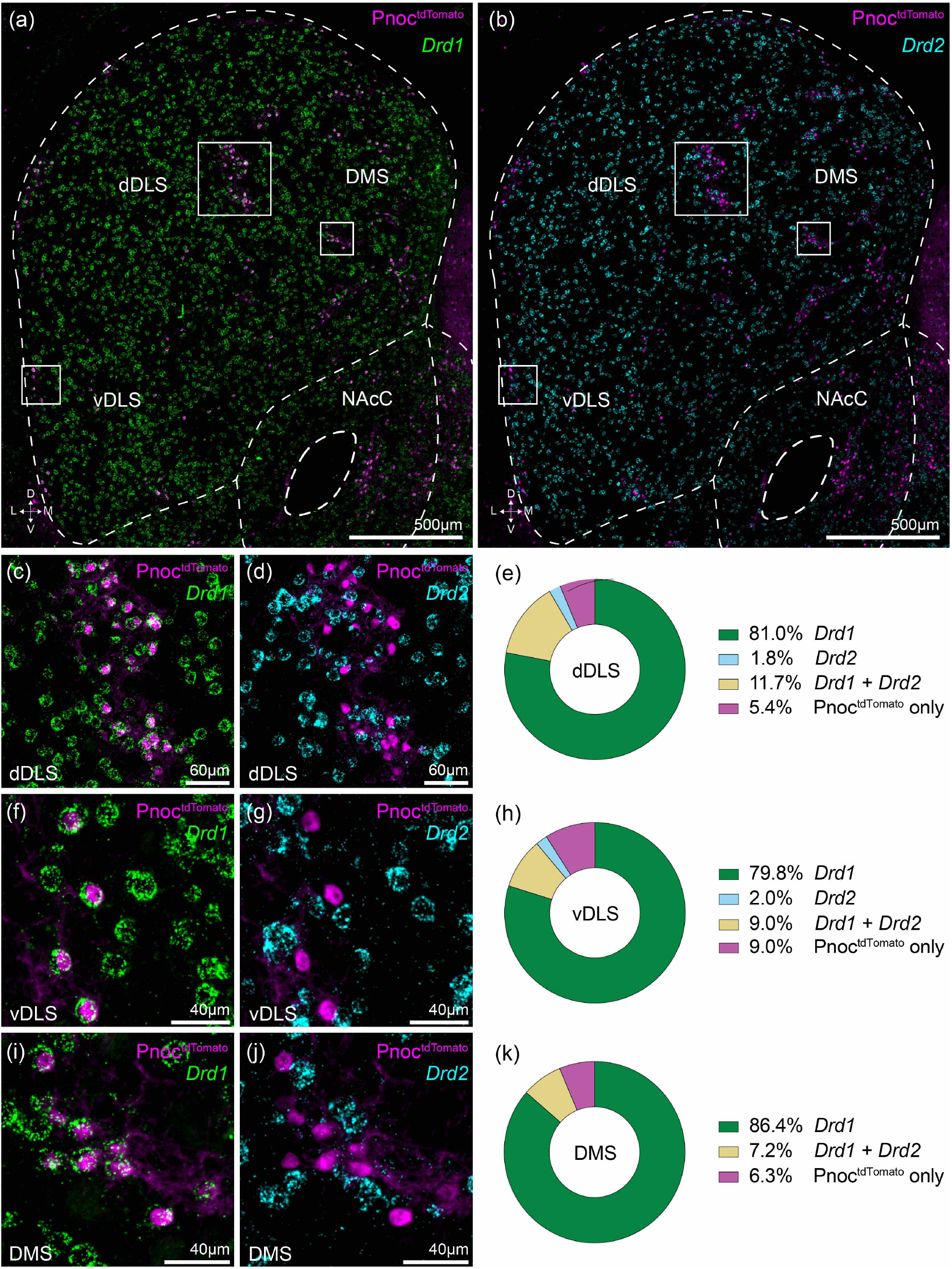
Pnoc^tdTomato^ expression across dorsal striatal subregions marks largely *Drd1* striatal neurons. Pnoc^tdTomato^ cells (magenta, IHC) co-labeled by ISH with (a) *Drd1* expression (green, ISH) or (b) *Drd2* expression (cyan, ISH) in the dorsal striatum. Higher magnification of insets from (a) and (b) in dorsal DLS (c,d), ventral DLS (f,g), and DMS (i,j). Summary of quantified Pnoc^tdTomato^ cells co-labeled by ISH with SPNs in dDLS (e), vDLS (h), and DMSl (k). dDLS = dorsal aspect of dorsolateral striatum; vDLS = ventral aspect of dorsolateral striatum; DMS = dorsomedial striatum.

**SUPPLEMENTAL FIGURE 2. - 2.**
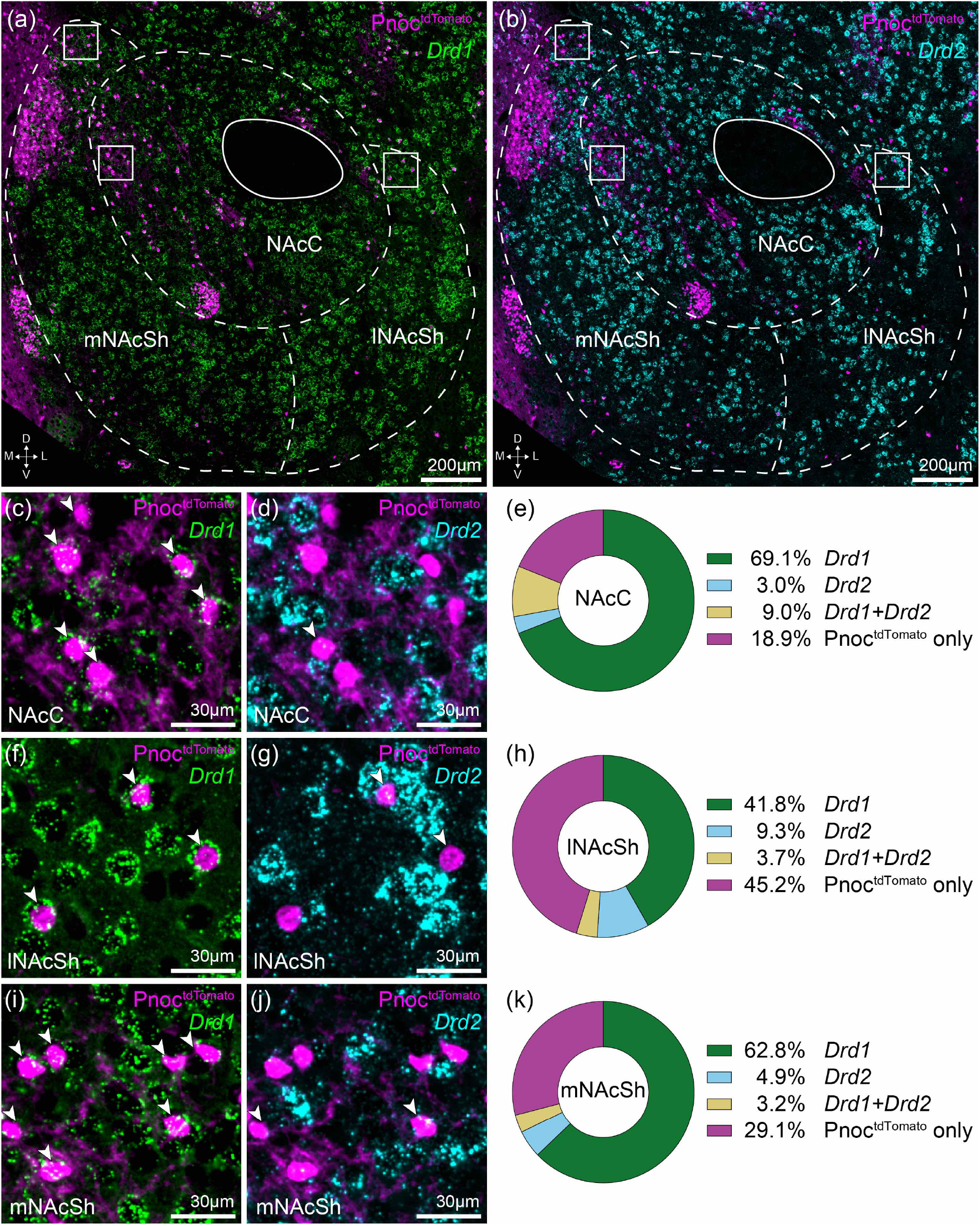
Pnoc^tdTomato^ expression in *Drd1* and *Drd2* striatal neurons varies across the nucleus accumbens. Pnoc^tdTomato^ cells (magenta, IHC) co-labeled by ISH with (a) *Drd1* expression (green, ISH) or (b) *Drd2* expression (cyan, ISH) in the nucleus accumbens. Higher magnification of insets from (a) and (b) in nucleus accumbens core (c,d), lateral shell (f,g), and medial shell (i,j). Summary of quantified Pnoc^tdTomato^ cells co-labeled by ISH with *Drd1* and/or *Drd2*-type SPNs in nucleus accumbens core (e), lateral shell (h), and medial shell (k). NAcC = nucleus accumbens core; lNAcSh = lateral accumbens shell; mNAcSh = medial accumbens shell.

**SUPPLEMENTAL FIGURE 3.**
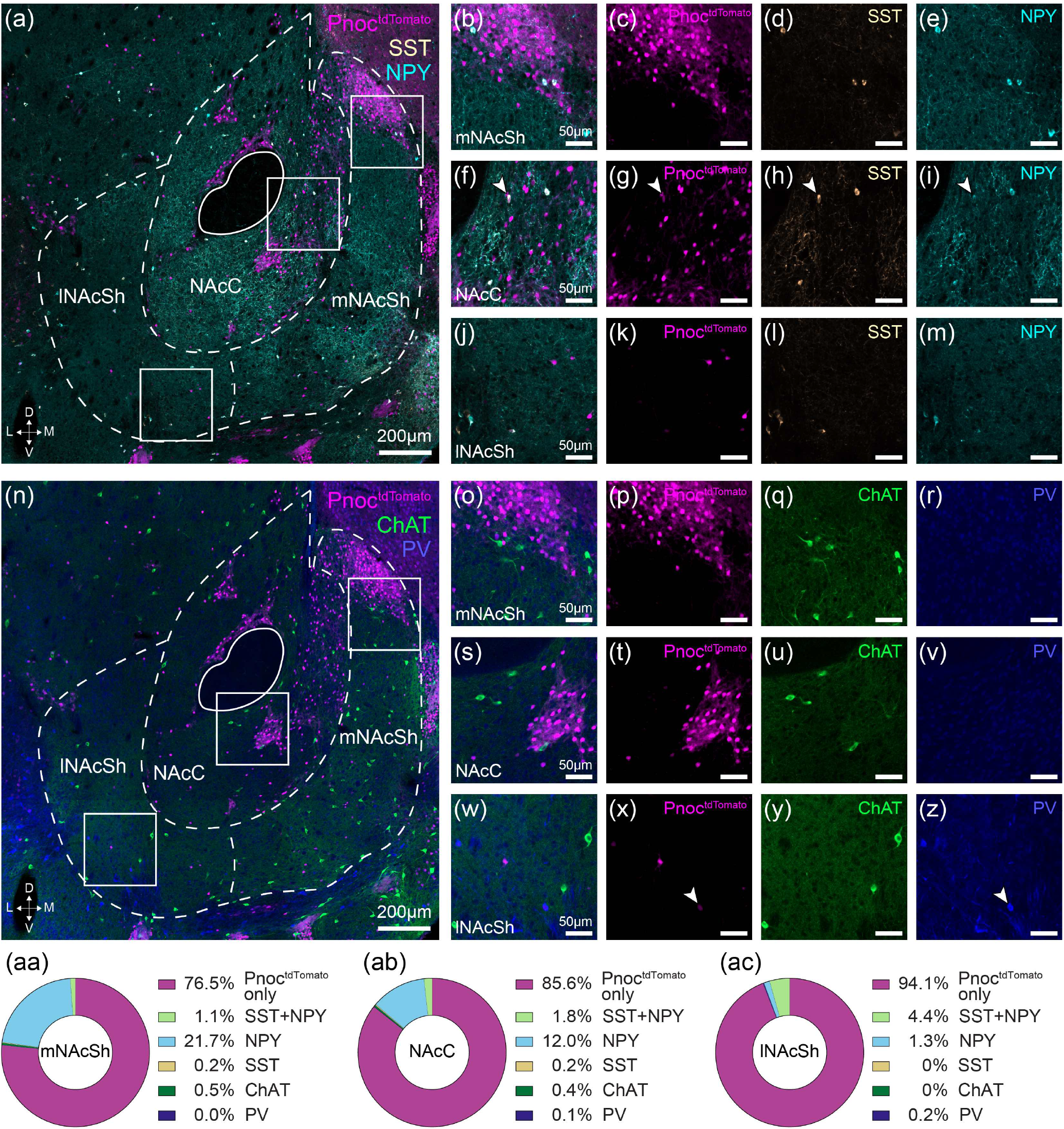
Pnoc^tdTomato^ NPY-labeled interneurons are enriched in the nucleus accumbens core and medial shell. (a) Pnoc^tdTomato^ cells (magenta, IHC) imaged with interneuron markers somatostatin (yellow, SST) and neuropeptide Y (cyan, NPY) in the nucleus accumbens. Higher magnification of insets from (a) in (b-e) medial accumbens shell, (f-i) accumbens core, and (j-m) lateral accumbens shell. (n-z) Same as (a-m) but for Pnoc^tdTomato^ cell colocalization with the interneuron markers choline acetyltransferase (green, ChAT) and parvalbumin (dark blue, PV) in the nucleus accumbens. (aa-ac) Summary of Pnoc^tdTomato^ colocalization with different interneuron markers in nucleus accumbens medial shell (aa), core (ab), and lateral shell (ac). NAcC = nucleus accumbens core; lNAcSh = lateral accumbens shell; mNAcSh = medial accumbens shell.

**SUPPLEMENTAL FIGURE 4. - 1.**
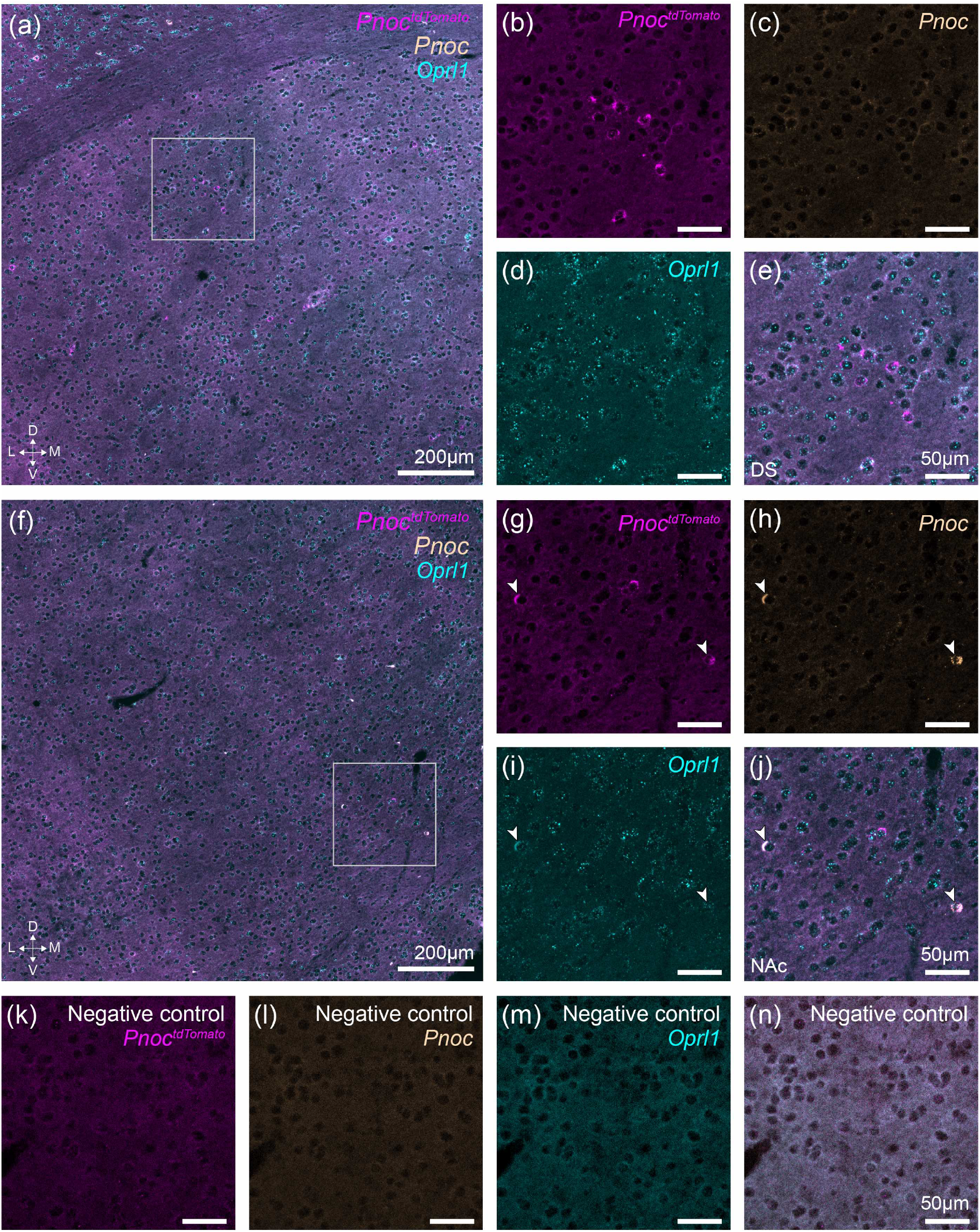
Low adult expression of Pnoc^tdTomato^ in the dorsal striatum does not recapitulate striosomal patterning of reporter. (a) Representative image showing ISH evaluation of Pnoc^tdTomato^ (*tdTomato*, magenta), *Pnoc* (yellow), and N/OFQ receptor gene *Oprl1* (cyan) expression in the dorsal striatum. (b-e) Higher magnification images of inset from (a) showing striosomal *tdTomato* expression that does not co-label with *Pnoc*, despite presence of *Oprl1*. (f) Representative image showing *tdTomato* (magenta), *Pnoc* (yellow), and *Oprl1* (cyan) ISH expression in the ventral striatum. (g-j) Higher magnification images of inset from (f) showing non-striosomal *tdTomato* expression that co-labels with *Pnoc* and *Oprl1*. (k-n) Negative control ISH probes. DS = dorsal striatum; NAc = nucleus accumbens.

**SUPPLEMENTAL FIGURE 4. - 2.**
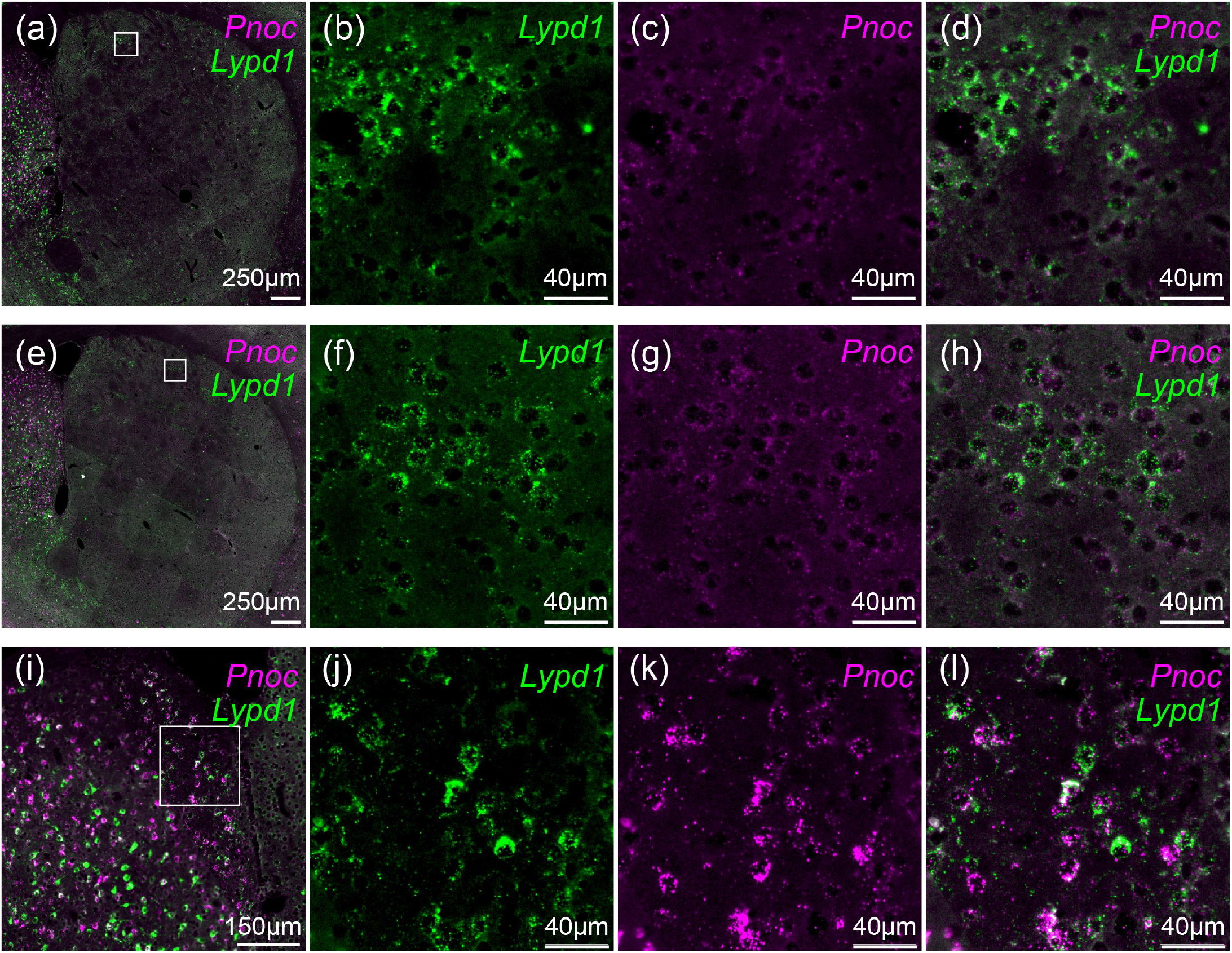
Adult chronic stress did not affect adult striosomal *Pnoc* expression. Striatal tissue collected after mice were exposed to no stress (a-d) or chronic immobilization stress (e-f; 1 hour/day for 14 days) for evaluation of adult dorsal striatal *Pnoc* mRNA expression (magenta) in striosomal regions (striosomal Lypd1, green). Neighboring septal region (i-l) expresses *Pnoc* robustly in adult tissue and serves as a positive control for *Pnoc* ISH labeling.

**SUPPLEMENTAL FIGURE 4. - 3.**
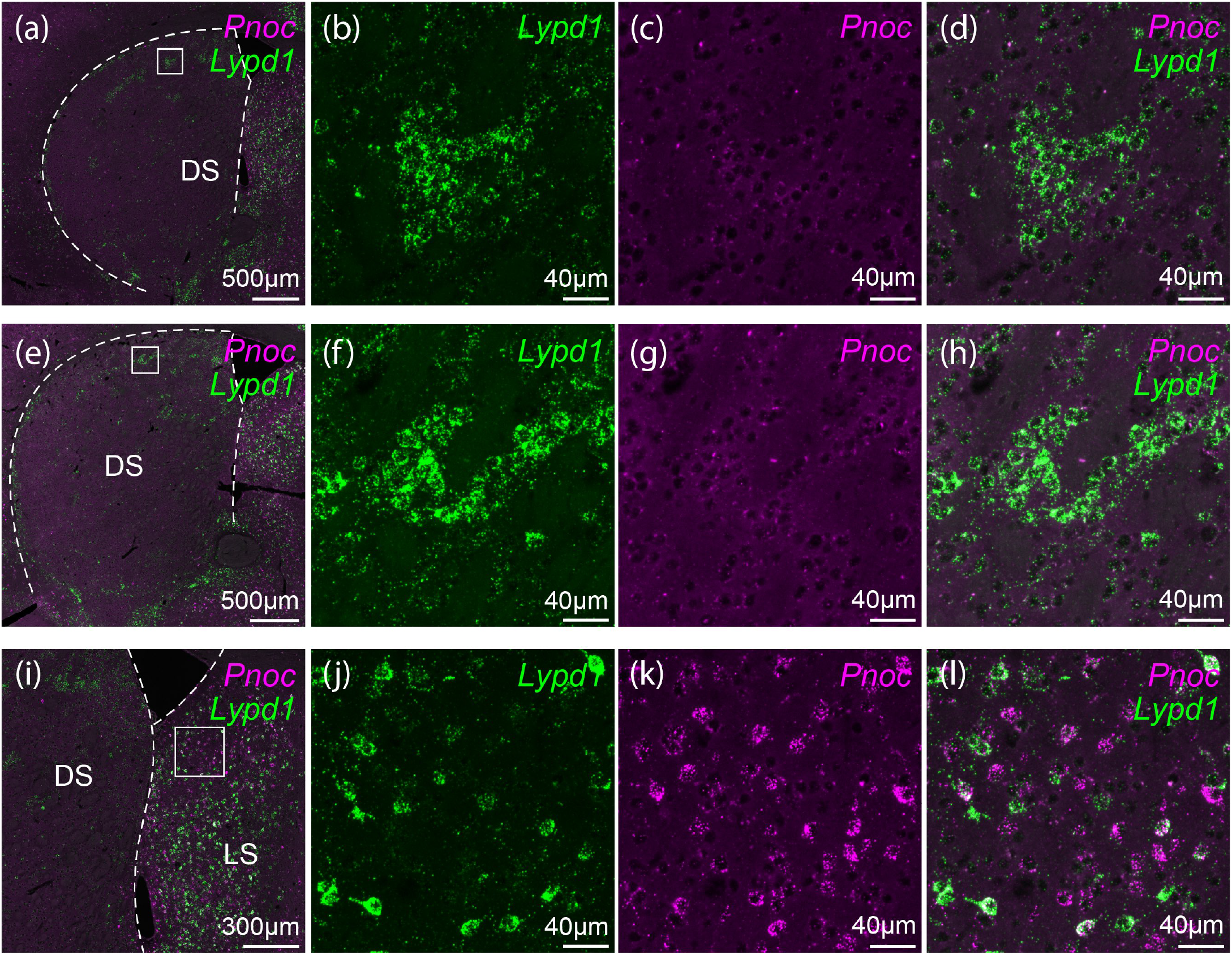
Adult exposure to chronic amphetamine does not detectably induce adult striosomal *Pnoc* mRNA expression. Striatal tissue collected after chronic exposure to saline (a-d) or amphetamine administration (e-h) was evaluated in striosomal regions (a, e; Lypd1, green; inset in b, f) for adult *Pnoc* mRNA expression in dorsal striatum (magenta, c, g); composite shown in d and h. Positive control for both *Lypd1* (j) and strong *Pnoc* signal (k) shown in neighboring septal tissue (i-l).

**SUPPLEMENTAL FIGURE 5.**
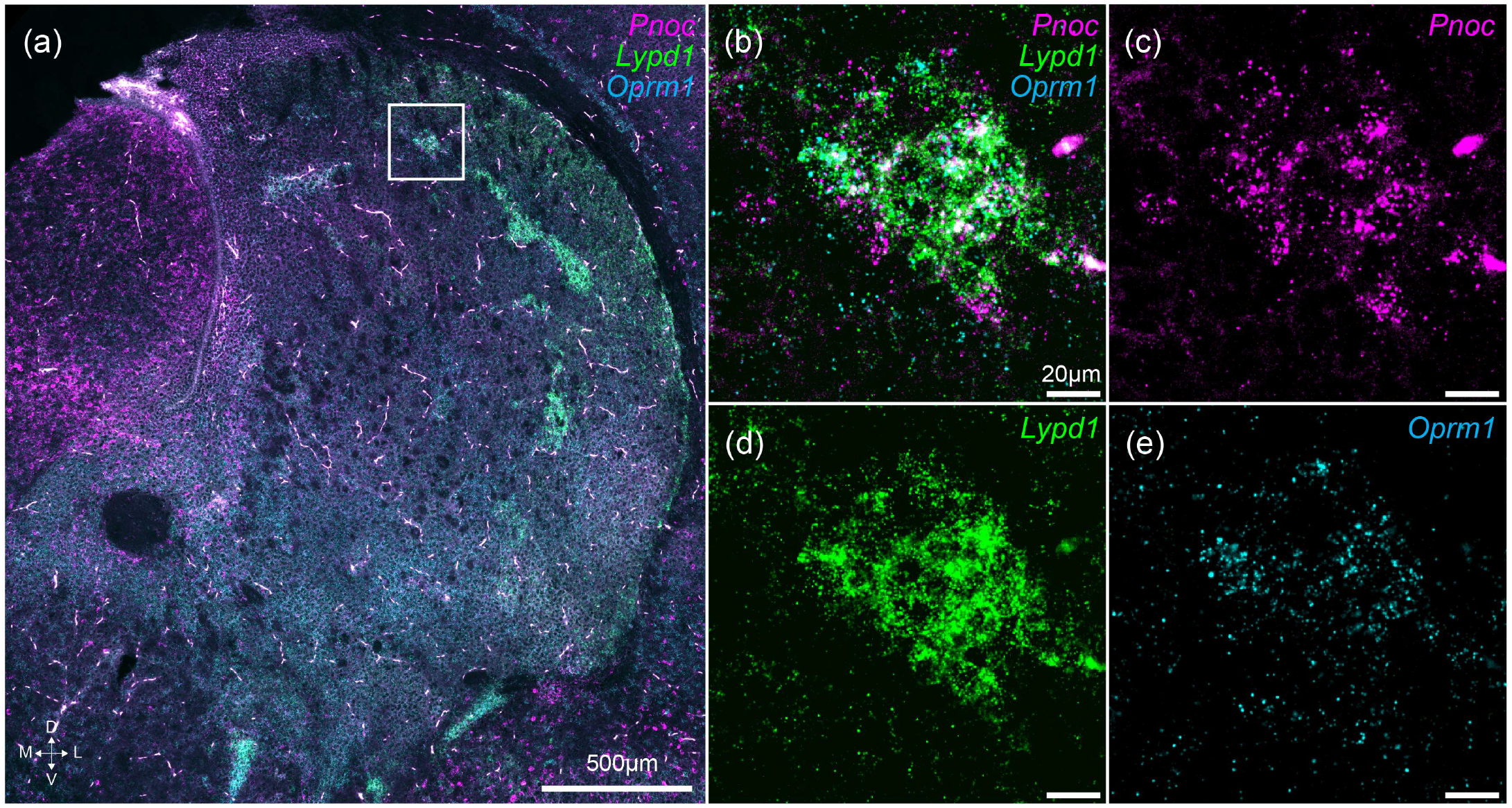
*Pnoc* expression exhibits striosomal patterning in neonate. (a) Striatal section of P5 neonatal tissue shows pattern of *Pnoc* mRNA (magenta) (b-c) overlapping striosomal marker expression (d) *Lypd1* (green) and (e) *Oprm1* (cyan).

## REFERENCES

Ahn, H.J., Hernandez, C.M., Levenson, J.M., Lubin, F.D., Liou, H.C. & Sweatt, J.D. (2008) c-Rel, an NF-κB family transcription factor, is required for hippocampal long-term synaptic plasticity and memory formation. Learning & Memory, 15(7), 539–549. doi:10.1101/lm.866408. URL https://www.ncbi.nlm.nih.gov/pmc/articles/PMC2505322/

Al-Hasani, R., McCall, J.G., Shin, G., Gomez, A.M., Schmitz, G.P., Bernardi, J.M. et al. (2015) Distinct Subpopulations of Nucleus Accumbens Dynorphin Neurons Drive Aversion and Reward. Neuron, 87(5), 1063. doi:10.1016/j.neuron.2015.08.019, publisher: NIH Public Access. URL https://www.ncbi.nlm.nih.gov/pmc/articles/PMC4625385/

Amemori, K.I., Amemori, S., Gibson, D.J. & Graybiel, A.M. (2018) Striatal Microstimulation Induces Persistent and Repetitive Negative Decision-Making Predicted by Striatal Beta-Band Oscillation. Neuron, 99(4), 829–841.e6. doi:10.1016/j.neuron.2018.07.022.

Amemori, K.i. & Graybiel, A.M. (2012) Localized microstimulation of primate pregenual cingulate cortex induces negative decision-making. Nature Neuroscience, 15(5), 776–785. doi:10.1038/nn.3088.

Amemori, S., Amemori, K.I., Yoshida, T., Papageorgiou, G.K., Xu, R., Shimazu, H. et al. (2020) Microstimulation of primate neocortex targeting striosomes induces negative decisionmaking. The European Journal of Neuroscience, 51(3), 731–741. doi:10.1111/ejn.14555.

Amemori, S., Graybiel, A.M. & Amemori, K.I. (2021) Causal Evidence for Induction of Pessimistic Decision-Making in Primates by the Network of Frontal Cortex and Striosomes. Frontiers in Neuroscience, 15, 649167. doi:10.3389/fnins.2021.649167.

Banghart, M.R., Neufeld, S.Q., Wong, N.C. & Sabatini, B.L. (2015) Enkephalin Disinhibits Mu Opioid Receptor-Rich Striatal Patches via Delta Opioid Receptors. Neuron, 88(6), 1227–1239.

Bankhead, P., Loughrey, M.B., Fernández, J.A., Dombrowski, Y., McArt, D.G., Dunne, P.D. et al. (2017) QuPath: Open source software for digital pathology image analysis. Scientific Reports, 7(1), 16878. doi:10.1038/s41598-017-17204-5.

Bebawy, D., Marquez, P., Samboul, S., Parikh, D., Hamid, A. & Lutfy, K. (2010) Orphanin FQ/nociceptin not only blocks but also reverses behavioral adaptive changes induced by repeated cocaine in mice. Biological Psychiatry, 68(3), 223–230. doi:10.1016/j.biopsych.2010.02.010.

Bloem, B., Huda, R., Amemori, K.I., Abate, A.S., Krishna, G., Wilson, A.L. et al. (2022) Multiplexed action-outcome representation by striatal striosome-matrix compartments detected with a mouse cost-benefit foraging task. Nature Communications, 13(1), 1541. doi:10.1038/s41467-022-28983-5.

Bloem, B., Huda, R., Sur, M. & Graybiel, A.M. (2017) Two-photon imaging in mice shows striosomes and matrix have overlapping but differential reinforcement-related responses. eLife, 6, e32353. doi:10.7554/eLife.32353.

Bodnar, R.J., Glass, M.J., Ragnauth, A. & Cooper, M.L. (1995) General, mu and kappa opioid antagonists in the nucleus accumbens alter food intake under deprivation, glucoprivic and palatable conditions. Brain Research, 700(1-2), 205–212. doi:10.1016/0006-8993(95)00957-r.

Boulanger, L.M. (2009) Immune proteins in brain development and synaptic plasticity. Neuron, 64(1), 93–109. doi:10.1016/j.neuron.2009.09.001.

Brimblecombe, K.R. & Cragg, S.J. (2017) The Striosome and Matrix Compartments of the Striatum: A Path through the Labyrinth from Neurochemistry toward Function. ACS Chem. Neurosci., 8(2), 235–242.

Canales, J.J. & Graybiel, A.M. (2000) A measure of striatal function predicts motor stereotypy. Nat. Neurosci., 3(4), 377–383.

Caputi, F.F., Di Benedetto, M., Carretta, D., Bastias del Carmen Candia, S., D’Addario, C., Cavina, C. et al. (2014) Dynorphin/KOP and nociceptin/NOP gene expression and epigenetic changes by cocaine in rat striatum and nucleus accumbens. Progress in Neuro-Psychopharmacology & Biological Psychiatry, 49, 36–46. doi:10.1016/j.pnpbp.2013.10.016.

Castro, D.C. & Berridge, K.C. (2014) Opioid Hedonic Hotspot in Nucleus Accumbens Shell: Mu, Delta, and Kappa Maps for Enhancement of Sweetness “Liking” and “Wanting”. The Journal of Neuroscience, 34(12), 4239–4250. doi:10.1523/JNEUROSCI.4458-13.2014. URL https://www.ncbi.nlm.nih.gov/pmc/articles/PMC3960467/

Castro, D.C. & Bruchas, M.R. (2019) A Motivational and Neuropeptidergic Hub: Anatomical and Functional Diversity within Nucleus Accumbens Shell. Neuron, 102(3), 529–552. doi:10.1016/j.neuron.2019.03.003. URL https://www.ncbi.nlm.nih.gov/pmc/articles/PMC6528838/

Castro, D.C., Oswell, C.S., Zhang, E.T., Pedersen, C.E., Piantadosi, S.C., Rossi, M.A. et al. (2021) An endogenous opioid circuit determines state-dependent reward consumption. Nature, 598(7882), 646–651. doi:10.1038/s41586-021-04013-0. URL https://www.ncbi.nlm.nih.gov/pmc/articles/PMC8858443/

Clark, S.D. & Abi-Dargham, A. (2019) The Role of Dynorphin and the Kappa Opioid Receptor in the Symptomatology of Schizophrenia: A Review of the Evidence. Biol. Psychiatry, 86(7), 502–511.

Crittenden, J.R. & Graybiel, A.M. (2011) Basal Ganglia disorders associated with imbalances in the striatal striosome and matrix compartments. Frontiers in Neuroanatomy, 5, 59. doi:10.3389/fnana.2011.00059.

Crittenden, J.R., Lacey, C.J., Weng, F.J., Garrison, C.E., Gibson, D.J., Lin, Y. et al. (2017) Striatal Cholinergic Interneurons Modulate Spike-Timing in Striosomes and Matrix by an Amphetamine-Sensitive Mechanism. Frontiers in Neuroanatomy, 11, 20. doi:10.3389/fnana.2017.00020.

Crittenden, J.R., Tillberg, P.W., Riad, M.H., Shima, Y., Gerfen, C.R., Curry, J. et al. (2016) Striosome-dendron bouquets highlight a unique striatonigral circuit targeting dopamine-containing neurons. Proceedings of the National Academy of Sciences of the United States of America, 113(40), 11318–11323. doi:10.1073/pnas.1613337113.

Cui, Y., Ostlund, S.B., James, A., Park, C.S., Ge, W., Roberts, K.W. et al. (2014) Targeted Expression of Mu-Opioid Receptors in a Subset of Striatal Direct-Pathway Neurons Restores Opiate Reward. Nature neuroscience, 17(2), 254–261. doi:10.1038/nn.3622. URL https://www.ncbi.nlm.nih.gov/pmc/articles/PMC4008330/

Cui, Y., Ostlund, S.B., James, A.S., Park, C.S., Ge, W., Roberts, K.W. et al. (2014) Targeted expression of μ-opioid receptors in a subset of striatal direct-pathway neurons restores opiate reward. Nat. Neurosci., 17(2), 254–261.

da Silva, J.A., Tecuapetla, F., Paixão, V. & Costa, R.M. (2018) Dopamine neuron activity before action initiation gates and invigorates future movements. Nature, 554(7691), 244–248. doi:10.1038/nature25457.

Damier, P., Hirsch, E.C., Agid, Y. & Graybiel, A.M. (1999) The substantia nigra of the human brain. II. Patterns of loss of dopamine-containing neurons in Parkinson’s disease. Brain: A Journal of Neurology, 122 (Pt 8), 1437–1448. doi:10.1093/brain/122.8.1437.

Der-Avakian, A., D’Souza, M.S., Potter, D.N., Chartoff, E.H., Carlezon, W.A., Pizzagalli, D.A. et al. (2017) Social defeat disrupts reward learning and potentiates striatal nociceptin/orphanin FQ mRNA in rats. Psychopharmacology, 234(9-10), 1603–1614. doi:10.1007/s00213-017-4584-y.

D’Oliveira da Silva, F., Robert, C., Lardant, E., Pizzano, C., Bruchas, M.R., Guiard, B.P. et al. (2023) Targeting Nociceptin/Orphanin FQ receptor to rescue cognitive symptoms in a mouse neuroendocrine model of chronic stress. Molecular Psychiatry,.doi:10.1038/s41380-023-02363-x.

Donica, C.L., Ramirez, V.I., Awwad, H.O., Zaveri, N.T., Toll, L. & Standifer, K.M. (2011) Orphanin FQ/nociceptin activates nuclear factor kappa B. Journal of Neuroimmune Pharmacology: The Official Journal of the Society on NeuroImmune Pharmacology, 6(4), 617–625. doi:10.1007/s11481-011-9279-2.

Eblen, F. & Graybiel, A.M. (1995) Highly restricted origin of prefrontal cortical inputs to striosomes in the macaque monkey. The Journal of Neuroscience: The Official Journal of the Society for Neuroscience, 15(9), 5999–6013. doi:10.1523/JNEUROSCI.15-09-05999.1995.

Enard, W. (2011) FOXP2 and the role of cortico-basal ganglia circuits in speech and language evolution. Current Opinion in Neurobiology, 21(3), 415–424. doi:10.1016/j.conb.2011.04.008.

Evans, R.C., Twedell, E.L., Zhu, M., Ascencio, J., Zhang, R. & Khaliq, Z.M. (2020) Functional Dissection of Basal Ganglia Inhibitory Inputs onto Substantia Nigra Dopaminergic Neurons. Cell Reports, 32(11), 108156. doi:10.1016/j.celrep.2020.108156.

Flau, K., Redmer, A., Liedtke, S., Kathmann, M. & Schlicker, E. (2002) Inhibition of striatal and retinal dopamine release via nociceptin/orphanin FQ receptors. British Journal of Pharmacology, 137(8), 1355–1361. doi:10.1038/sj.bjp.0704998.

Fong, W.L., Kuo, H.Y., Wu, H.L., Chen, S.Y. & Liu, F.C. (2018) Differential and Overlapping Pattern of Foxp1 and Foxp2 Expression in the Striatum of Adult Mouse Brain. Neuroscience, 388, 214–223. doi:10.1016/j.neuroscience.2018.07.017.

Friedman, A., Homma, D., Bloem, B., Gibb, L.G., Amemori, K.I., Hu, D. et al. (2017) Chronic Stress Alters Striosome-Circuit Dynamics, Leading to Aberrant Decision-Making. Cell, 171(5), 1191–1205.e28. doi:10.1016/j.cell.2017.10.017.

Friedman, A., Homma, D., Gibb, L.G., Amemori, K.I., Rubin, S.J., Hood, A.S. et al. (2015) A Corticostriatal Path Targeting Striosomes Controls Decision-Making under Conflict. Cell, 161(6), 1320–1333. doi:10.1016/j.cell.2015.04.049.

Friedman, A., Hueske, E., Drammis, S.M., Toro Arana, S.E., Nelson, E.D., Carter, C.W. et al. (2020) Striosomes Mediate Value-Based Learning Vulnerable in Age and a Huntington’s Disease Model. Cell, 183(4), 918–934.e49. doi:10.1016/j.cell.2020.09.060.

Fu, X., Zhu, Z.H., Wang, Y.Q. & Wu, G.C. (2007) Regulation of proinflammatory cytokines gene expression by nociceptin/orphanin FQ in the spinal cord and the cultured astrocytes. Neuroscience, 144(1), 275–285. doi:10.1016/j.neuroscience.2006.09.016.

Gavioli, E.C. & Calo’, G. (2013) Nociceptin/orphanin FQ receptor antagonists as innovative antidepressant drugs. Pharmacology & Therapeutics, 140(1), 10–25. doi:10.1016/j.pharmthera.2013.05.008.

Giguère, N., Burke Nanni, S. & Trudeau, L.E. (2018) On Cell Loss and Selective Vulnerability of Neuronal Populations in Parkinson’s Disease. Frontiers in Neurology, 9, 455. doi:10.3389/fneur.2018.00455.

Gokce, O., Stanley, G.M., Treutlein, B., Neff, N.F., Camp, J.G., Malenka, R.C. et al. (2016) Cellular Taxonomy of the Mouse Striatum as Revealed by Single-Cell RNA-Seq. Cell Reports, 16(4), 1126–1137. doi:10.1016/j.celrep.2016.06.059.

Graybiel, A.M. (1990) Neurotransmitters and neuromodulators in the basal ganglia. Trends Neurosci., 13(7), 244–254.

Graybiel, A.M. & Hickey, T.L. (1982) Chemospecificity of ontogenetic units in the striatum: demonstration by combining [3H]thymidine neuronography and histochemical staining. Proceedings of the National Academy of Sciences of the United States of America, 79(1), 198–202. doi:10.1073/pnas.79.1.198.

Graybiel, A.M. & Matsushima, A. (2023) Striosomes and Matrisomes: Scaffolds for Dynamic Coupling of Volition and Action. Annual Review of Neuroscience, 46, 359–380. doi:10.1146/annurev-neuro-121522-025740.

Hardaway, J.A., Halladay, L.R., Mazzone, C.M., Pati, D., Bloodgood, D.W., Kim, M. et al. (2019) Central Amygdala Prepronociceptin-Expressing Neurons Mediate Palatable Food Consumption and Reward. Neuron, 102(5), 1088. doi:10.1016/j.neuron.2019.04.036.

He, M., Tucciarone, J., Lee, S., Nigro, M.J., Kim, Y., Levine, J.M. et al. (2016) Strategies and Tools for Combinatorial Targeting of GABAergic Neurons in Mouse Cerebral Cortex. Neuron, 92(2), 555. doi:10.1016/j.neuron.2016.10.009.

Hernandez, J., Perez, L., Soto, R., Le, N., Gastelum, C. & Wagner, E.J. (2021) Nociceptin/orphanin FQ neurons in the Arcuate Nucleus and Ventral Tegmental Area Act via Nociceptin Opioid Peptide Receptor Signaling to Inhibit Proopiomelanocortin and A10 Dopamine Neurons and Thereby Modulate Ingestion of Palatable Food. Physiology & Behavior, 228, 113183. doi:10.1016/j.physbeh.2020.113183.

Hikosaka, O., Kim, H.F., Yasuda, M. & Yamamoto, S. (2014) Basal ganglia circuits for reward value-guided behavior. Annual Review of Neuroscience, 37, 289–306. doi:10.1146/annurev-neuro-071013-013924.

Howe, M.W. & Dombeck, D.A. (2016) Rapid signalling in distinct dopaminergic axons during locomotion and reward. Nature, 535(7613), 505–510. doi:10.1038/nature18942.

Imperato, A., Angelucci, L., Casolini, P., Zocchi, A. & Puglisi-Allegra, S. (1992) Repeated stressful experiences differently affect limbic dopamine release during and following stress. Brain Research, 577(2), 194–199. doi:10.1016/0006-8993(92)90274-d.

Ironside, M., Amemori, K.I., McGrath, C.L., Pedersen, M.L., Kang, M.S., Amemori, S. et al. (2020) Approach-Avoidance Conflict in Major Depressive Disorder: Congruent Neural Findings in Humans and Nonhuman Primates. Biological Psychiatry, 87(5), 399–408. doi:10.1016/j.biopsych.2019.08.022.

Jenck, F., Moreau, J.L., Martin, J.R., Kilpatrick, G.J., Reinscheid, R.K., Monsma, F.J. et al. (1997) Orphanin FQ acts as an anxiolytic to attenuate behavioral responses to stress. Proceedings of the National Academy of Sciences of the United States of America, 94(26), 14854–14858. doi:10.1073/pnas.94.26.14854.

Kash, T.L., Pleil, K.E., Marcinkiewcz, C.A., Lowery-Gionta, E.G., Crowley, N., Mazzone, C. et al. (2015) Neuropeptide regulation of signaling and behavior in the BNST. Molecules and Cells, 38(1), 1–13. doi:10.14348/molcells.2015.2261.

Kelly, S.M., Raudales, R., He, M., Lee, J.H., Kim, Y., Gibb, L.G. et al. (2018) Radial Glial Lineage Progression and Differential Intermediate Progenitor Amplification Underlie Striatal Compartments and Circuit Organization. Neuron, 99(2), 345–361.e4. doi:10.1016/j.neuron.2018.06.021.

Kennedy, S.E., Koeppe, R.A., Young, E.A. & Zubieta, J.K. (2006) Dysregulation of Endogenous Opioid Emotion Regulation Circuitry in Major Depression in Women. Arch. Gen. Psychiatry, 63(11), 1199–1208. Publisher: American Medical Association.

Koizumi, M., Midorikawa, N., Takeshima, H. & Murphy, N.P. (2004) Exogenous, but not endogenous nociceptin modulates mesolimbic dopamine release in mice. Journal of Neurochemistry, 89(1), 257–263. doi:10.1111/j.1471-4159.2003.02322.x.

Koob, G.F. (2008) A role for brain stress systems in addiction. Neuron, 59(1), 11–34. doi:10.1016/j.neuron.2008.06.012.

Kordower, J.H., Olanow, C.W., Dodiya, H.B., Chu, Y., Beach, T.G., Adler, C.H. et al. (2013) Disease duration and the integrity of the nigrostriatal system in Parkinson’s disease. Brain: A Journal of Neurology, 136(Pt 8), 2419–2431. doi:10.1093/brain/awt192.

Lambert, D.G. (2008) The nociceptin/orphanin FQ receptor: a target with broad therapeutic potential. Nature Reviews. Drug Discovery, 7(8), 694–710. doi:10.1038/nrd2572.

Lança, A.J., Boyd, S., Kolb, B.E. & van der Kooy, D. (1986) The development of a patchy organization of the rat striatum. Brain Research, 392(1-2), 1–10. doi:10.1016/0165-3806(86)90226-9.

Lee, I.B., Lee, E., Han, N.E., Slavuj, M., Hwang, J.W., Lee, A. et al. (2024) Persistent enhancement of basolateral amygdala-dorsomedial striatum synapses causes compulsive-like behaviors in mice. Nature Communications, 15(1), 219. doi:10.1038/s41467-023-44322-8.

Lutfy, K., Do, T. & Maidment, N.T. (2001) Orphanin FQ/nociceptin attenuates motor stimulation and changes in nucleus accumbens extracellular dopamine induced by cocaine in rats. Psychopharmacology, 154(1), 1–7. doi:10.1007/s002130000609.

Lutfy, K., Khaliq, I., Carroll, F.I. & Maidment, N.T. (2002) Orphanin FQ/nociceptin blocks cocaine-induced behavioral sensitization in rats. Psychopharmacology, 164(2), 168–176. doi:10.1007/s00213-002-1192-1.

Madisen, L., Zwingman, T.A., Sunkin, S.M., Oh, S.W., Zariwala, H.A., Gu, H. et al. (2010) A robust and high-throughput Cre reporting and characterization system for the whole mouse brain. Nature Neuroscience, 13(1), 133–140. doi:10.1038/nn.2467.

Mangiavacchi, S., Masi, F., Scheggi, S., Leggio, B., De Montis, M.G. & Gambarana, C. (2001) Long-term behavioral and neurochemical effects of chronic stress exposure in rats. Journal of Neurochemistry, 79(6), 1113–1121. doi:10.1046/j.1471-4159.2001.00665.x.

Margolis, E.B., Moulton, M.G., Lambeth, P.S. & O’Meara, M.J. (2023) The life and times of endogenous opioid peptides: Updated understanding of synthesis, spatiotemporal dynamics, and the clinical impact in alcohol use disorder. Neuropharmacology, 225, 109376.

Marquez, P., Nguyen, A.T., Hamid, A. & Lutfy, K. (2008) The endogenous OFQ/N/ORL-1 receptor system regulates the rewarding effects of acute cocaine. Neuropharmacology, 54(3), 564–568. doi:10.1016/j.neuropharm.2007.11.003.

Marti, M., Mela, F., Veronesi, C., Guerrini, R., Salvadori, S., Federici, M. et al. (2004) Blockade of nociceptin/orphanin FQ receptor signaling in rat substantia nigra pars reticulata stimulates nigrostriatal dopaminergic transmission and motor behavior. The Journal of Neuroscience: The Official Journal of the Society for Neuroscience, 24(30), 6659–6666. doi:10.1523/JNEUROSCI.0987-04.2004.

Matsushima, A. & Graybiel, A.M. (2020) Combinatorial Developmental Controls on Striatonigral Circuits. Cell Reports, 31(11), 107778. doi:10.1016/j.celrep.2020.107778.

Medeiros, I.U., Ruzza, C., Asth, L., Guerrini, R., Romão, P.R.T., Gavioli, E.C. et al. (2015) Blockade of nociceptin/orphanin FQ receptor signaling reverses LPS-induced depressive-like behavior in mice. Peptides, 72, 95–103. doi:10.1016/j.peptides.2015.05.006.

Menalled, L. & Brunner, D. (2014) Animal models of Huntington’s disease for translation to the clinic: best practices. Movement Disorders: Official Journal of the Movement Disorder Society, 29(11), 1375–1390. doi:10.1002/mds.26006.

Mendez, I., Sanchez-Pernaute, R., Cooper, O., Viñuela, A., Ferrari, D., Björklund, L. et al. (2005) Cell type analysis of functional fetal dopamine cell suspension transplants in the striatum and substantia nigra of patients with Parkinson’s disease. Brain: A Journal of Neurology, 128(Pt 7), 1498–1510. doi:10.1093/brain/awh510.

Meunier, J.C., Mollereau, C., Toll, L., Suaudeau, C., Moisand, C., Alvinerie, P. et al. (1995) Isolation and structure of the endogenous agonist of opioid receptor-like ORL1 receptor. Nature, 377(6549), 532–535. doi:10.1038/377532a0.

Mogil, J.S., Grisel, J.E., Reinscheid, R.K., Civelli, O., Belknap, J.K. & Grandy, D.K. (1996) Orphanin FQ is a functional anti-opioid peptide. Neuroscience, 75(2), 333–337. doi:10.1016/0306-4522(96)00338-7.

Mollereau, C. & Mouledous, L. (2000) Tissue distribution of the opioid receptor-like (ORL1) receptor. Peptides, 21(7), 907–917. doi:10.1016/s0196-9781(00)00227-8.

Morigaki, R., Lee, J.H., Yoshida, T., Wüthrich, C., Hu, D., Crittenden, J.R. et al. (2020) Spatiotemporal Up-Regulation of Mu Opioid Receptor 1 in Striatum of Mouse Model of Huntington’s Disease Differentially Affecting Caudal and Striosomal Regions. Front. Neuroanat., 14, 608060.

Murphy, N.P. & Maidment, N.T. (1999) Orphanin FQ/nociceptin modulation of mesolimbic dopamine transmission determined by microdialysis. Journal of Neurochemistry, 73(1), 179–186. doi:10.1046/j.1471-4159.1999.0730179.x.

Märtin, A., Calvigioni, D., Tzortzi, O., Fuzik, J., Wärnberg, E. & Meletis, K. (2019) A Spatiomolecular Map of the Striatum. Cell Reports, 29(13), 4320–4333.e5. doi:10.1016/j.celrep.2019.11.096.

Neal, C.R., Mansour, A., Reinscheid, R., Nothacker, H.P., Civelli, O. & Watson, S.J. (1999) Localization of orphanin FQ (nociceptin) peptide and messenger RNA in the central nervous system of the rat. The Journal of Comparative Neurology, 406(4), 503–547.

Norton, C.S., Neal, C.R., Kumar, S., Akil, H. & Watson, S.J. (2002) Nociceptin/orphanin FQ and opioid receptor-like receptor mRNA expression in dopamine systems. The Journal of Comparative Neurology, 444(4), 358–368. doi:10.1002/cne.10154.

Olianas, M.C., Dedoni, S., Boi, M. & Onali, P. (2008) Activation of nociceptin/orphanin FQ-NOP receptor system inhibits tyrosine hydroxylase phosphorylation, dopamine synthesis, and dopamine D(1) receptor signaling in rat nucleus accumbens and dorsal striatum. Journal of Neurochemistry, 107(2), 544–556. doi:10.1111/j.1471-4159.2008.05629.x.

Parker, K.E., Pedersen, C.E., Gomez, A.M., Spangler, S.M., Walicki, M.C., Feng, S.Y. et al. (2019) A Paranigral VTA Nociceptin Circuit that Constrains Motivation for Reward. Cell, 178(3), 653–671.e19. doi:10.1016/j.cell.2019.06.034.

Pedersen, M.L., Ironside, M., Amemori, K.I., McGrath, C.L., Kang, M.S., Graybiel, A.M. et al. (2021) Computational phenotyping of brain-behavior dynamics underlying approach-avoidance conflict in major depressive disorder. PLoS computational biology, 17(5), e1008955. doi:10.1371/journal.pcbi.1008955.

Pereira Luppi, M., Azcorra, M., Caronia-Brown, G., Poulin, J.F., Gaertner, Z., Gatica, S. et al. (2021) Sox6 expression distinguishes dorsally and ventrally biased dopamine neurons in the substantia nigra with distinctive properties and embryonic origins. Cell Reports, 37(6), 109975. doi:10.1016/j.celrep.2021.109975.

Post, A., Smart, T.S., Krikke-Workel, J., Dawson, G.R., Harmer, C.J., Browning, M. et al. (2016) A Selective Nociceptin Receptor Antagonist to Treat Depression: Evidence from Preclinical and Clinical Studies. Neuropsychopharmacology: Official Publication of the American College of Neuropsychopharmacology, 41(7), 1803–1812. doi:10.1038/npp.2015.348.

Ragnauth, A., Moroz, M. & Bodnar, R.J. (2000) Multiple opioid receptors mediate feeding elicited by mu and delta opioid receptor subtype agonists in the nucleus accumbens shell in rats. Brain Research, 876(1-2), 76–87. doi:10.1016/s0006-8993(00)02631-7.

Ragsdale, C.W. & Graybiel, A.M. (1988) Fibers from the basolateral nucleus of the amygdala selectively innervate striosomes in the caudate nucleus of the cat. The Journal of Comparative Neurology, 269(4), 506–522. doi:10.1002/cne.902690404.

Reinscheid, R.K., Nothacker, H.P., Bourson, A., Ardati, A., Henningsen, R.A., Bunzow, J.R. et al. (1995) Orphanin FQ: a neuropeptide that activates an opioidlike G protein-coupled receptor. Science (New York, N.Y.), 270(5237), 792–794. doi:10.1126/science.270.5237.792.

Resendez, S.L., Dome, M., Gormley, G., Franco, D., Nevárez, N., Hamid, A.A. et al. (2013) μ-Opioid Receptors within Subregions of the Striatum Mediate Pair Bond Formation through Parallel Yet Distinct Reward Mechanisms. The Journal of Neuroscience, 33(21), 9140–9149. doi:10.1523/JNEUROSCI.4123-12.2013. URL https://www.ncbi.nlm.nih.gov/pmc/articles/PMC6705037/

Rolle, C.E., Pedersen, M.L., Johnson, N., Amemori, K.I., Ironside, M., Graybiel, A.M. et al. (2022) The Role of the Dorsal-Lateral Prefrontal Cortex in Reward Sensitivity During Approach-Avoidance Conflict. Cerebral Cortex (New York, N.Y.: 1991), 32(6), 1269–1285. doi:10.1093/cercor/bhab292.

Romualdi, P., Di Benedetto, M., D’Addario, C., Collins, S.L., Wade, D., Candeletti, S. et al. (2007) Chronic cocaine produces decreases in N/OFQ peptide levels in select rat brain regions. Journal of molecular neuroscience: MN, 31(2), 159–164. doi:10.1385/jmn/31:02:159.

Saito, Y., Maruyama, K., Kawano, H., Hagino-Yamagishi, K., Kawamura, K., Saido, T.C. et al. (1996) Molecular cloning and characterization of a novel form of neuropeptide gene as a developmentally regulated molecule. The Journal of Biological Chemistry, 271(26), 15615–15622. doi:10.1074/jbc.271.26.15615.

Saito, Y., Maruyama, K., Saido, T.C. & Kawashima, S. (1995) N23K, a gene transiently up-regulated during neural differentiation, encodes a precursor protein for a newly identified neuropeptide nociceptin. Biochemical and Biophysical Research Communications, 217(2), 539–545. doi:10.1006/bbrc.1995.2809.

Saito, Y., Maruyama, K., Saido, T.C. & Kawashima, S. (1997) Overexpression of a neuropeptide nociceptin/orphanin FQ precursor gene, N23K/N27K, induces neurite outgrowth in mouse NS20Y cells. Journal of Neuroscience Research, 48(5), 397–406. doi:10.1002/(sici)1097-4547(19970601)48:5<397::aid-jnr2>3.0.co;2-9.

Sakoori, K. & Murphy, N.P. (2004) Central administration of nociceptin/orphanin FQ blocks the acquisition of conditioned place preference to morphine and cocaine, but not conditioned place aversion to naloxone in mice. Psychopharmacology, 172(2), 129–136. doi:10.1007/s00213-003-1643-3.

Saunders, A., Macosko, E.Z., Wysoker, A., Goldman, M., Krienen, F.M., de Rivera, H. et al. (2018) Molecular Diversity and Specializations among the Cells of the Adult Mouse Brain. Cell, 174(4), 1015–1030.e16. doi:10.1016/j.cell.2018.07.028.

Schultz, W. (2016) Dopamine reward prediction-error signalling: a two-component response. Nature Reviews. Neuroscience, 17(3), 183–195. doi:10.1038/nrn.2015.26.

Sgroi, S. & Tonini, R. (2018) Opioidergic Modulation of Striatal Circuits, Implications in Parkinson’s Disease and Levodopa Induced Dyskinesia. Front. Neurol., 9, 524.

Smith, J.B., Klug, J.R., Ross, D.L., Howard, C.D., Hollon, N.G., Ko, V.I. et al. (2016) Genetic-Based Dissection Unveils the Inputs and Outputs of Striatal Patch and Matrix Compartments. Neuron, 91(5), 1069–1084. doi:10.1016/j.neuron.2016.07.046.

Smith, M.A., Choudhury, A.I., Glegola, J.A., Viskaitis, P., Irvine, E.E., de Campos Silva, P.C.C. et al. (2020) Extrahypothalamic GABAergic nociceptin-expressing neurons regulate AgRP neuron activity to control feeding behavior. The Journal of Clinical Investigation, 130(1), 126–142. doi:10.1172/JCI130340.

Tajima, K. & Fukuda, T. (2013) Region-specific diversity of striosomes in the mouse striatum revealed by the differential immunoreactivities for mu-opioid receptor, substance P, and enkephalin. Neuroscience, 241, 215–228.

Tejeda, H.A., Wu, J., Kornspun, A.R., Pignatelli, M., Kashtelyan, V., Krashes, M.J. et al. (2017) Pathway and Cell-Specific Kappa-Opioid Receptor Modulation of Excitatory-Inhibitory Balance Differentially Gates D1 and D2 Accumbens Neuron Activity. Neuron, 93(1), 147–163. doi:10.1016/j.neuron.2016.12.005. URL https://www.ncbi.nlm.nih.gov/pmc/articles/PMC5808882/

Ubaldi, M., Cannella, N., Borruto, A.M., Petrella, M., Micioni Di Bonaventura, M.V., Soverchia, L. et al. (2021) Role of Nociceptin/Orphanin FQ-NOP Receptor System in the Regulation of Stress-Related Disorders. International Journal of Molecular Sciences, 22(23), 12956. doi:10.3390/ijms222312956.

Vazquez-DeRose, J., Stauber, G., Khroyan, T.V., Xie, X.S., Zaveri, N.T. & Toll, L. (2013) Retrodialysis of N/OFQ into the nucleus accumbens shell blocks cocaine-induced increases in extracellular dopamine and locomotor activity. European Journal of Pharmacology, 699(1-3), 200–206. doi:10.1016/j.ejphar.2012.11.050.

Watabe-Uchida, M., Zhu, L., Ogawa, S.K., Vamanrao, A. & Uchida, N. (2012) Whole-brain mapping of direct inputs to midbrain dopamine neurons. Neuron, 74(5), 858–873. doi:10.1016/j.neuron.2012.03.017.

Wilson, L., Klausner, M., Chuang, S., Patel, S. & Pratt, W.E. (2024) An examination of the effects of nucleus accumbens core nociceptin on appetitive and consummatory motivation for food. Behavioural Brain Research, 462, 114895. doi:10.1016/j.bbr.2024.114895.

Witkin, J.M., Wallace, T.L. & Martin, W.J. (2019) Therapeutic Approaches for NOP Receptor Antagonists in Neurobehavioral Disorders: Clinical Studies in Major Depressive Disorder and Alcohol Use Disorder with BTRX-246040 (LY2940094). Handbook of Experimental Pharmacology, 254, 399–415. doi:10.1007/164_2018_186.

Yamada, T., McGeer, P.L., Baimbridge, K.G. & McGeer, E.G. (1990) Relative sparing in Parkinson’s disease of substantia nigra dopamine neurons containing calbindin-D28K. Brain Research, 526(2), 303–307. doi:10.1016/0006-8993(90)91236-a.

Yoshizawa, T., Ito, M. & Doya, K. (2018) Reward-Predictive Neural Activities in Striatal Striosome Compartments. eNeuro, 5(1), ENEURO.0367–17.2018. doi:10.1523/ENEURO.0367-17.2018.

Zaveri, N.T. (2011) The nociceptin/orphanin FQ receptor (NOP) as a target for drug abuse medications. Current Topics in Medicinal Chemistry, 11(9), 1151–1156. doi:10.2174/156802611795371341.

Zaveri, N.T., Waleh, N. & Toll, L. (2006) Regulation of the prepronociceptin gene and its effect on neuronal differentiation. Gene, 384, 27–36. doi:10.1016/j.gene.2006.07.007.

Zheng, F., Grandy, D.K. & Johnson, S.W. (2002) Actions of orphanin FQ/nociceptin on rat ventral tegmental area neurons in vitro. British Journal of Pharmacology, 136(7), 1065–1071. doi:10.1038/sj.bjp.0704806.

